# PES-8 is required for Cytoskeletal Organization and Contractility in the *C. elegans* Spermatheca

**DOI:** 10.64898/2026.04.08.717217

**Authors:** Fereshteh Sadeghian, Erin J Cram

**Affiliations:** Bioengineering, Northeastern University, Boston, MA; Biology, Northeastern University, Boston, MA

## Abstract

Proper regulation of contraction and relaxation in biological tubes is essential for organismal function. In *C. elegans*, the spermatheca, composed of smooth muscle-like cells, undergoes repeated stretching and contraction as oocytes pass through. Here we describe PES-8, a previously uncharacterized protein, as a regulator of spermatheca contractility. PES-8 contains a predicted extracellular zona pellucida-like domain and an unstructured cytoplasmic tail, suggesting dual roles in extracellular and cytoplasmic signaling. PES-8 localizes to the plasma membrane of the spermatheca, the spermathecal-uterine valve, and uterus. Functional analysis shows that PES-8 is essential for spermathecal function; its loss disrupts actomyosin fiber alignment, FLN-1/filamin localization, apical junction organization, and Ca²⁺ signaling, preventing oocyte transit. These findings identify PES-8 as a key regulator of cytoskeletal organization and calcium-mediated contractility in the *C. elegans* spermatheca.

## Introduction

The actomyosin cytoskeleton plays essential roles in cell migration (Weißenbruch & Mayor, 2024), tissue development (Miao & Blankenship, 2020), wound repair (Hui et al., 2022), and the mechanical adaptation of cells and tissues to stress (Ennomani et al., 2016; Sadeghian et al., 2024). In smooth muscle cells, the anchoring and alignment of actomyosin regulates cell shape and contractility, contributing to the function of tissues found in the cardiovascular, respiratory and reproductive systems (Donadon & Santoro, 2021). Dysregulation of smooth muscle cell contractility can contribute to disorders such as asthma (Donadon & Santoro, 2021). Despite decades of work on the regulation of actin and myosin, many questions remain regarding how these dynamic actin networks generate and coordinate contractile force within living animals. To address this, we use *C. elegans* as a model to study actomyosin-driven contraction in their physiological context, where the mechanical and signaling environment is preserved.

In the *C. elegans* somatic gonad, the spermatheca, a single-layered tube comprising 24 smooth muscle-like cells, undergoes cyclic stretching, constriction, and relaxation as approximately 150 oocytes pass through (Figure 1). Immediately following oocyte entry, fertilization occurs and the spermatheca contracts, pushing the fertilized embryo into the uterus. Stretching the spermathecal cells activates phospholipase PLC-1/PLCε, leading to inositol trisphosphate (IP_3_) and diacylglycerol (DAG) generation. IP_3_ induces the release of calcium from the ER, which, coupled with RHO-1/Rho signaling and NMY-1/myosin activation, results in cell contraction in the spermathecal bag (Bouffard et al., 2019; Kelley et al., 2018). Disrupting any part of this pathway causes irregular calcium signaling and defective contractions (Bouffard et al., 2019; Castaneda et al., 2020; Kelley et al., 2018; Sadeghian et al., 2022).

**Figure 1:**
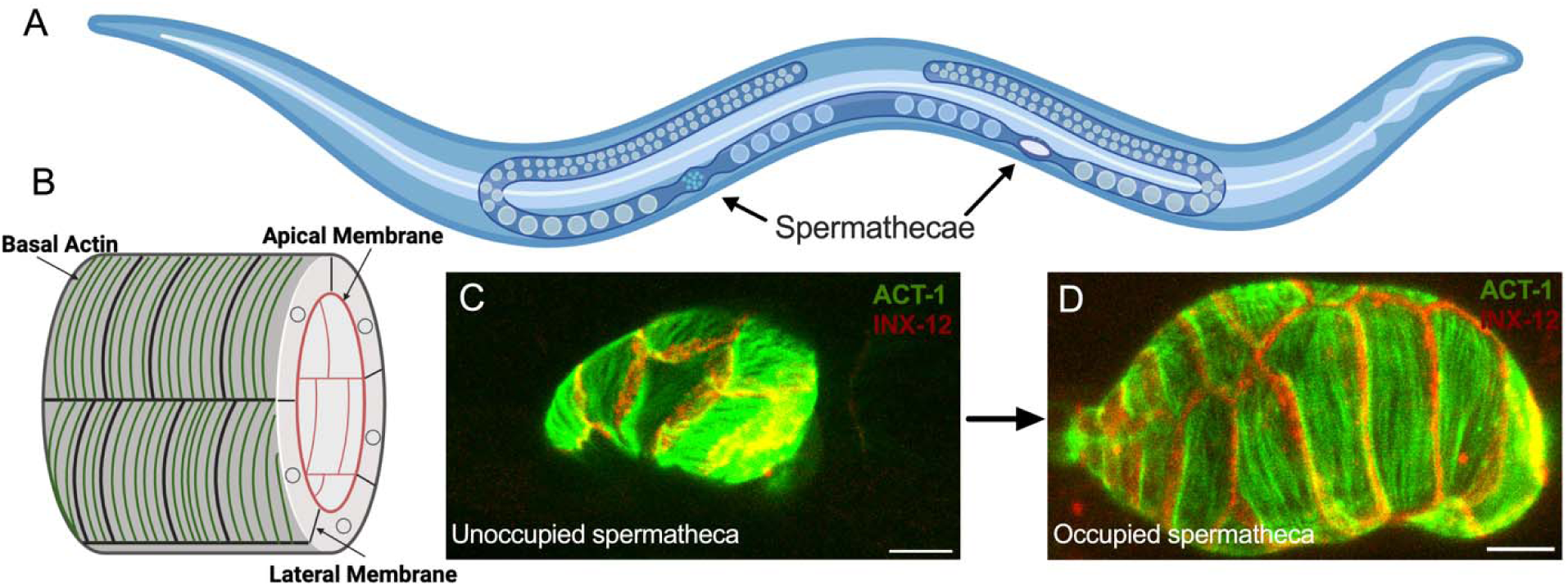
A) *C. elegans* anatomy. B) A cartoon of a tube representing apical, basal and lateral sides of the spermatheca. Spermatheca expressing labeled actin (ACT-1::GFP) and a gap junction marker (INX-12::mApple) to label the lateral cell membranes C) before ovulation and D) during ovulation through the spermatheca. Scale bar, 20 μm.

Contraction of the spermatheca is coordinated through the activation of NMY-1/non-muscle myosin, which generates the force needed to contract the actomyosin network (Kelley et al., 2018; Wirshing & Cram, 2017). This network consists of basal actomyosin bundles that are evenly spaced and aligned along the cell’s long axis (Strome, 1986; Wirshing & Cram, 2017). Before the first ovulation, the actin network is mesh-like and non-aligned; however, during the first ovulation, parallel fiber bundles assemble and remain organized throughout subsequent ovulations. Myosin activity is essential for establishing this ordered arrangement (Kelley et al., 2018; Wirshing & Cram, 2017). In addition to myosin, multiple other proteins contribute to the formation and maintenance of the organized actin cytoskeleton in the spermatheca, including filamin (Kovacevic & Cram, 2010), spectrin (Wirshing & Cram, 2018), actin-interacting protein 1 (Ono & Ono, 2014), polarity regulators (Aono et al., 2004), formin-related proteins (Hegsted et al., 2016), and a gelsolin-like protein (Deng et al., 2007).

Spermathecal cells are connected at the apical side by *C. elegans* apical junctions (CeAJs), which polarize epithelial cells, provide barrier function, and connect to the cytoskeleton to mediate mechanical signal transmission across tissues (Lynch, 2009). The CeAJ includes the cadherin–catenin complex (CCC), composed of HMR-1/E-cadherin, HMP-1/α-catenin, and HMP-2/β-catenin (Lynch, 2009). The CCC is required for ventral enclosure during embryogenesis and directly binds actin to anchor the cytoskeleton (Costa et al., 1998). Loss of either HMP-1/α-catenin or HMP-2/β-catenin disrupts this connection, causing actin bundles to detach from the CeAJ in embryos (Costa et al., 1998). Directly basal to the CCC is the DLG-1/Discs large and AJM-1/Apical junction molecules complex (DAC), which is in close proximity to the septate junctions (Lockwood et al., 2008; Lynch, 2009). DLG-1 is required for localization of the coiled-coil protein AJM-1 around the junctional belt (Köppen et al., 2001).

In a screen for regulators of spermathecal contractility, we identified the novel protein PES-8 (Sadeghian et al., 2025). PES-8 is a protein of unknown function that lacks sequence conservation outside of nematodes. Modeling with AlphaFold suggests PES-8 is a transmembrane protein with a ZP-like extracellular domain. The Zona Pellucida (ZP) domain is a protein-protein interaction domain involved in a wide range of functions, including sperm-egg fusion and as a structural element in apical extracellular matrix in the vulva, excretory canal and other structures in nematode larvae (Cohen et al., 2020). Typically, ZP-domain proteins polymerize into filamentous or matrix-like structures, and can serve as attachment points, for example, for neurite extension (Heiman & Bülow, 2024; Jovine et al., 2005). Here we show that when PES-8 is depleted by RNAi or lost through mutation, spermathecal actomyosin fiber alignment, FLN-1/filamin localization, apical junction organization, and Ca²⁺ signaling are all disrupted, leading to a failure of oocyte transit through the spermatheca and reduced fertility in *C. elegans*.

## Results

### PES-8 Structure

PES-8 is an uncharacterized protein of unknown function. The *pes-8* gene spans ∼3.5 kb and is composed of 11 exons (Figure 2A and B). The *pes-8* gene has two predicted isoforms, *pes-8a* and *pes-8b*, both of which are expressed (Figure S1 and S2). PES-8 has low sequence homology outside of nematodes. AlphaFold and SwissModel predict a C-terminal signal peptide (Figure S3A), an alpha-helical signal sequence, a transmembrane domain (Figure S3B) and a beta-sheet protein structure, as well as an unstructured cytoplasmic tail (Figure 2C-E) (Duckert et al., 2004). To identify possible functions for PES-8, we used Foldseek, an Alphafold-based algorithm that identifies proteins with similar folds (van Kempen et al., 2024). Foldseek identified similarity between the PES-8 beta sheet structure and the Zona Pellucida (ZP) domain, a bipartite immunoglobulin-like protein interaction domain (Monné et al., 2008). Domains that fold similarly are found in the ZP-domain proteins such as the cuticulins in *C. elegans* (Figure 2H), TGFβ receptor type 3 (Figure 2I), uromodulin (UMOD), the T-cell receptor delta variable 1 in humans, and other proteins (Table S1). Since the ZP-like domains in ZP3 and UMOD tend to polymerize (Bokhove & Jovine, 2018), we used AlphaFold to model multimers that may be formed by PES-8 (Figure S1 and S2). PES-8 is predicted to contain one immunoglobulin-like fold, unlike most ZP-domains, which are composed of two (Litscher & Wassarman, 2020). Most ZP-domain proteins are expressed as membrane tethered proteins, with the ZP domain released extracellularly by protease cleavage (Nishio et al., 2024). PES-8 is predicted to encode a transmembrane protein lacking a furin cleavage site, suggesting it will remain anchored to the cell membrane (Duckert et al., 2004).

**Figure 2:**
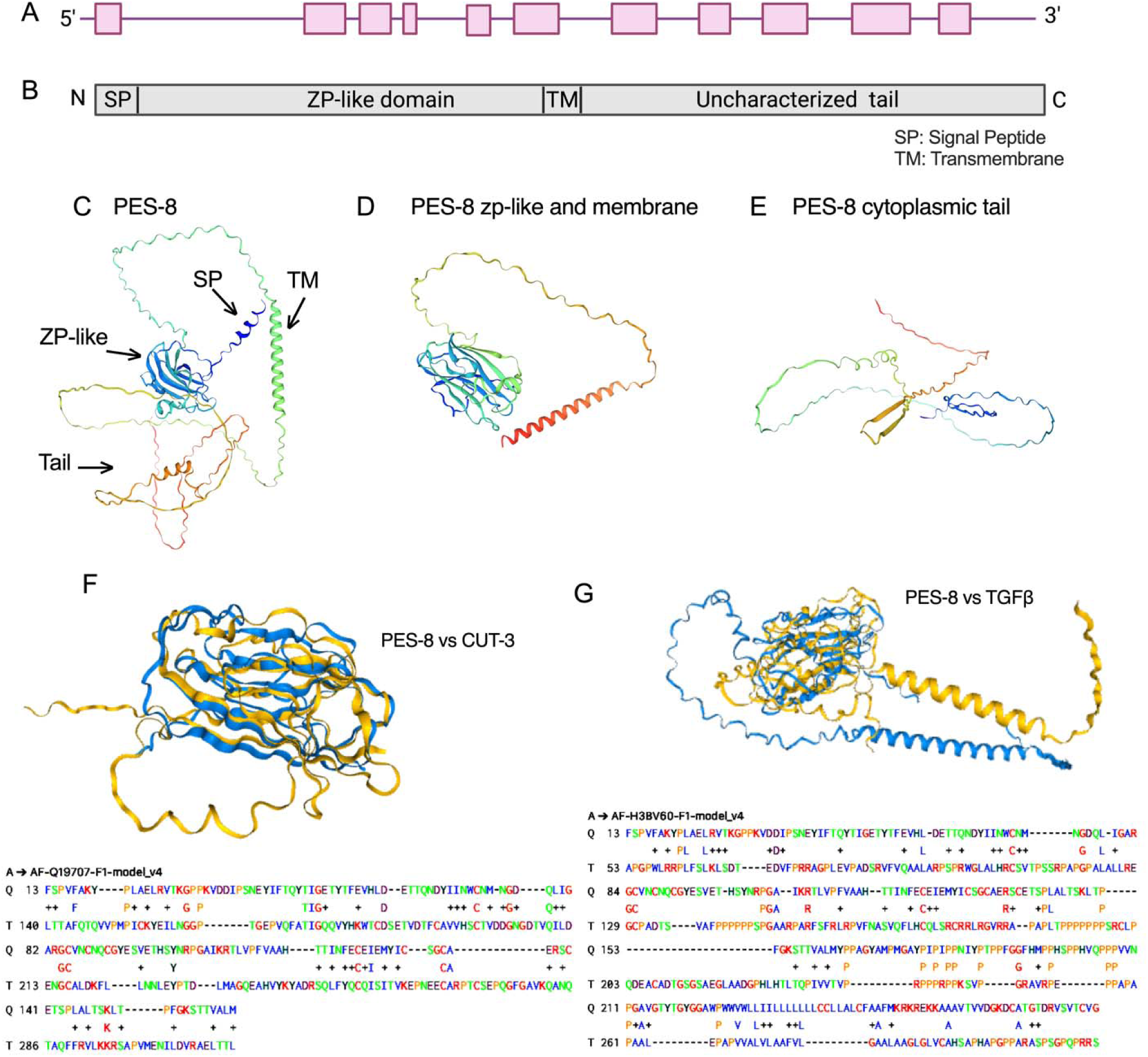
PES-8 structure. A) The *pes-8* gene spans ∼3.5 kb and is composed of 11 exons. B) PES-8 has a predicted signal peptide domain at the N-terminus, a ZP-like domain, a membrane domain in the middle, and an unstructured C-terminal tail. Models of the PES-8 (C), the PES-8 ZP-like domain and transmembrane domains (D), and the PES-8 cytoplasmic tail sequence (E) as predicted by SwissModel prediction tool (https://swissmodel.expasy.org/) (Schwede et al., 2003). F) PES-8 (blue, Q) and CUT-3 (yellow, T) are predicted to fold similarly. G) PES-8 (blue, Q) and a domain from the extracellular portion of TGFβ (yellow, T) are predicted to fold similarly. Sequence conservation is shown below the models (van Kempen et al., 2024).

### PES-8 localizes to the spermathecal membrane

To visualize PES-8 expression, we generated transgenic nematodes expressing *pes-8p*::PES-8::GFP. Because the N-terminal signal sequence is predicted to be cleaved, we placed GFP at the C-terminus of the *pes-8* genomic sequence and injected the construct into N2 worms. PES-8::GFP is expressed in spermathecal bag, sp-ut valve and uterus (Figure 3A-C). In the spermatheca, PES-8 is enriched on the apical and lateral membranes and is seen in punctae on the basal surface of the spermathecal bag (Figure 3D and E) (Movie S1).

**Figure 3:**
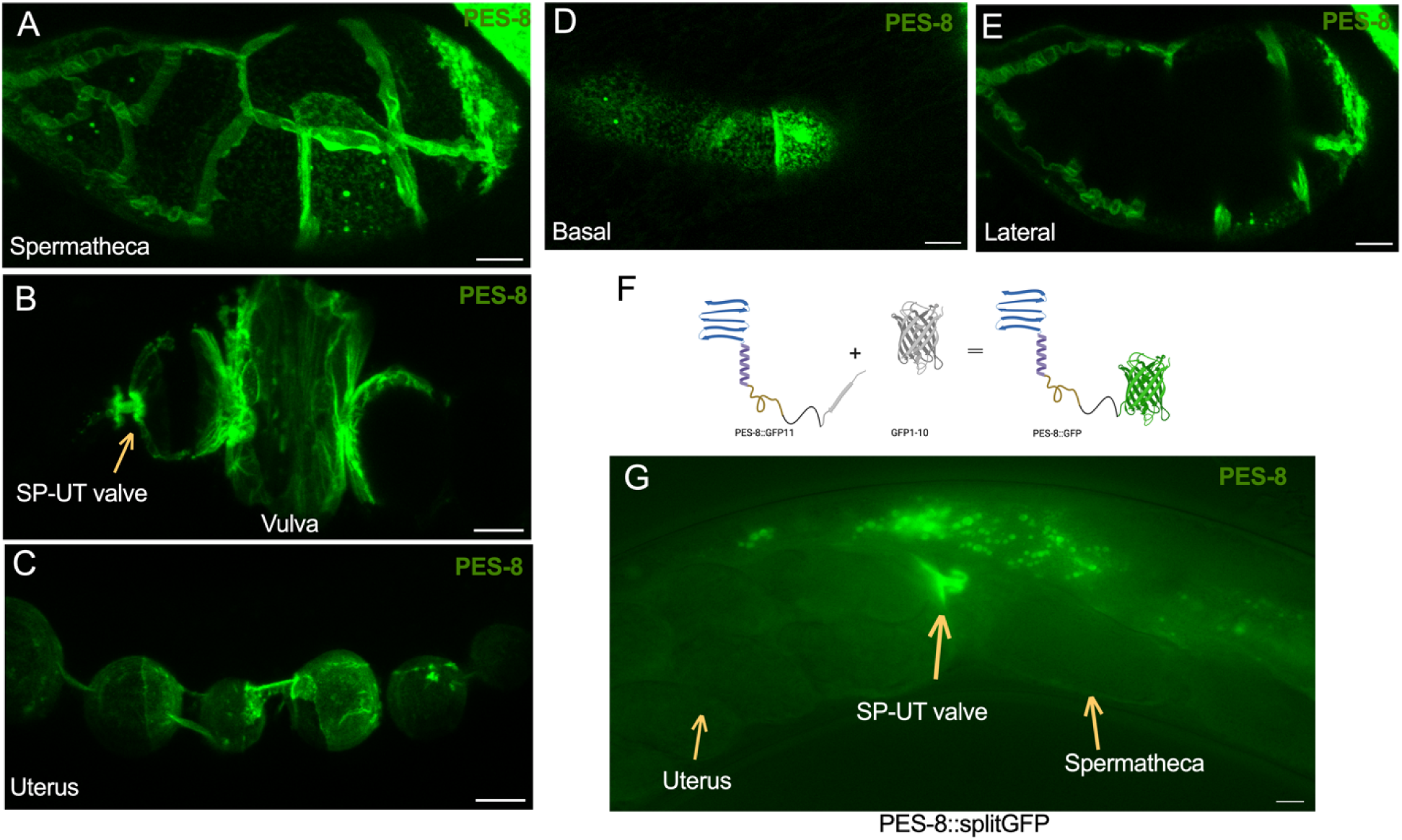
PES-8 expression in *C. elegans*. A-C) *pes-8*p::PES-8::GFP expressed in the spermatheca (A), sp-ut valve and vulva (B) and uterus (C). D and E) PES-8::GFP is expressed on the basal (D) and lateral (E) sides of spermatheca (Movie S1). F) Diagram of Split GFP fused with PES-8. G) Expression of PES-8::Split GFP in spermatheca, sp-ut valve, and uterus. Scale bar, 20 μm in A, D, E. Scale bar, 30 μm in B, C, G.

To determine the orientation of PES-8 in the cell membrane, we used Split GFP, an engineered tool which consists of two parts, GFP1-10 and GFP11, to identify the cytoplasmic domain of the protein. A separation between the 10th and 11th strands of the GFP beta barrel splits the large GFP1–10 from the small GFP11 fragment. These fragments are not fluorescent unless they bind to each other in the cell or tissue (Figure 3F) (Goudeau et al., 2021). We added GFP-11 to the C-terminus of PES-8 (*pes-8p*::PES-8::GFP11) and crossed the resulting transgenic strain with a strain broadly expressing cytoplasmic GFP1-10. Imaging of the crossed strain showed green membrane localization in the spermatheca, sp-ut valve, and uterus (Figure 3G). This result confirmed that the C-terminus of PES-8 is in the cytoplasm.

### PES-8 is required for oocyte transit in spermatheca

To study the function of PES-8, we generated three *pes-8* mutants (*xb7, xb13* and *xb14*) using CRISPR-Cas9 genome editing (Figure 4A). Two guide RNAs targeting sequences upstream and downstream of *pes-8* resulted in a complete gene deletion, *pes-8(xb7)*. Four additional guide RNAs targeted the cytoplasmic and extracellular sequences of the *pes-8* gene, to generate *xb13* and *xb14*, respectively. *xb13* preserves the signal sequence, extracellular domain, and transmembrane domain, and *xb14* preserves the signal sequence, transmembrane domain and cytoplasmic domain.

**Figure 4:**
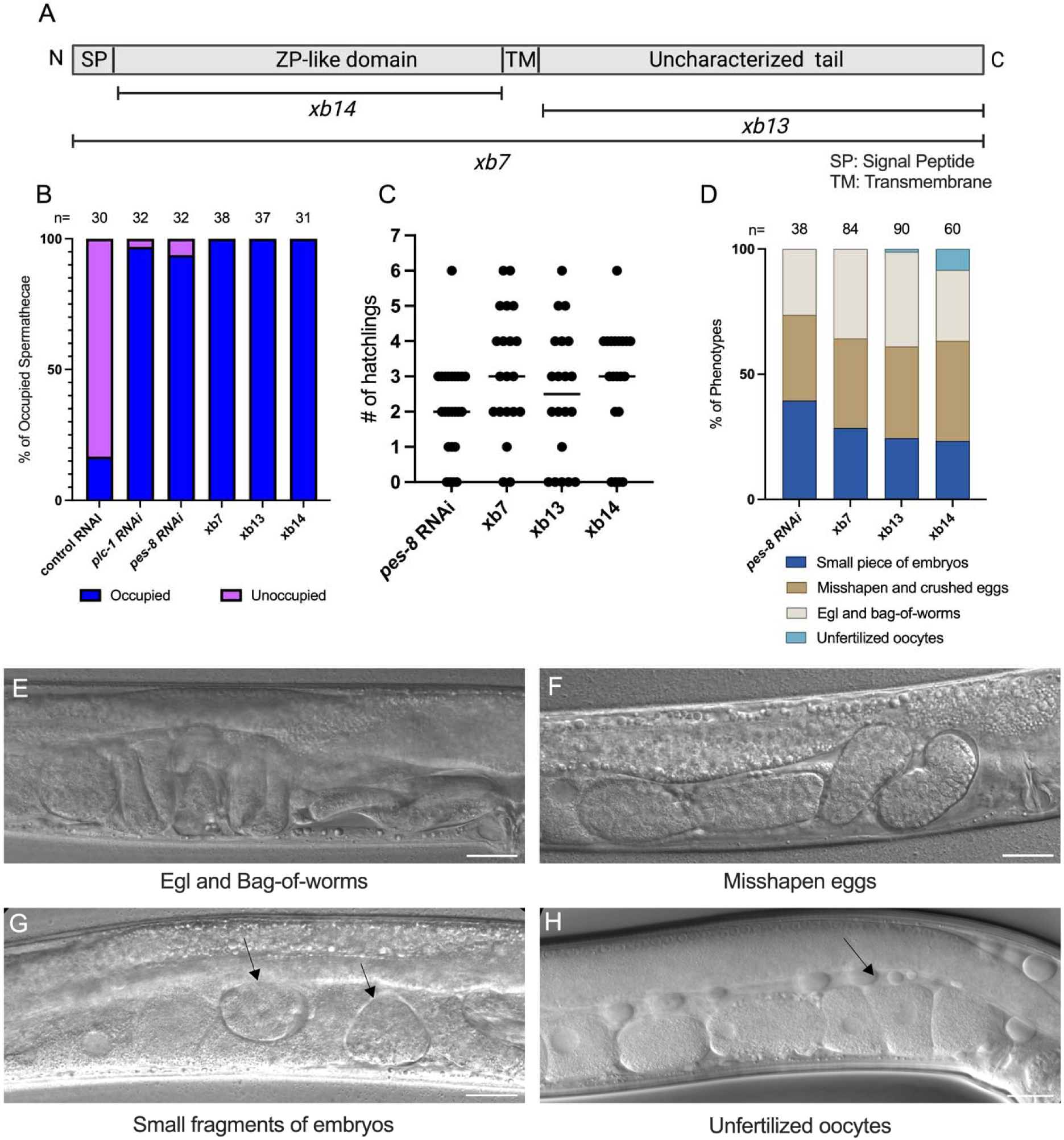
Loss of *pes-8* causes ovulation and fertility phenotypes. A) Diagrammatic representation of *pes-8* mutants including the entire *pes-8* deletion *xb7*, ZP-like domain deletion *xb14*, and the cytoplasmic tail deletion *xb13*. B) Percentages of spermathecae occupied by an oocyte in a GCaMP expressing strain with control (empty vector and *plc-1* RNAi), *pes-8* RNAi, and alleles are shown. C) Number of mutants and *pes-8* RNAi hatchlings, (n=20 for each). D) Phenotypes exhibited by *pes-8* RNAi and mutant alleles. Representative DIC images displaying E) the egg-laying defective (egl) and bag-of-worms phenotype, F) misshapen eggs, G) small fragments of embryos (indicated by the arrows), and H) unfertilized oocytes in the uterus (indicated by the arrows). Scale bar, 30 μm.

To examine the role of PES-8 in *C. elegans* spermatheca contractility, we scored spermathecal occupancy in these mutants and in a GCaMP-expressing strain treated with *pes-8* RNAi (Figure 4B). GCaMP was used to visualize the spermatheca with fluorescence illumination. In control animals, ∼20% of spermathecae were occupied by an oocyte. In all *pes-8* mutants, 100% of the spermathecae are occupied with an oocyte. In *pes-8* RNAi, 95% of spermathecae were occupied with an oocyte, similar to the effect of the positive control *plc-1* RNAi and phenocopying the mutant alleles. This suggests PES-8 is required for transit of oocytes through the spermatheca.

The mutant animals are also egg-laying defective (egl); embryos that do enter the uterus are not laid and hatch inside the mother, resulting in the bag-of-worms phenotype. Mutant adults typically produce only a few progeny (Figure 4C) and then die within a few days. We also observed crushed ooplasm, small eggs, and unfertilized eggs in the uterus for both RNAi knock-down and deletion animals (Figure 4 D-H), suggesting that PES-8 is essential for proper oocyte transit through the spermatheca, for egg laying, and for fertility.

### PES-8 is required for basal fiber anchorage and actomyosin bundle organization

Because contraction of the spermatheca is driven by actomyosin (Kelley & Cram, 2019), we next explored whether PES-8 was required for formation, alignment or anchorage of the basal actin fiber bundles in the spermatheca. To observe actin bundles in the *pes-8* mutants, we used phalloidin staining, since it was not possible to cross these animals to introduce the ACT-1::GFP transgene. Phalloidin staining revealed normal actin orientation in dissected wild-type spermathecae (Figure 5A). In *pes-8* mutants, the actin bundles were disrupted (Figure 5B-D). Similarly, *pes-8* RNAi in animals expressing ACT-1::GFP and H2B::mCherry led to disrupted actin bundles and aggregation of actin to the nuclei of the cells (Figure 5E and F).

**Figure 5:**
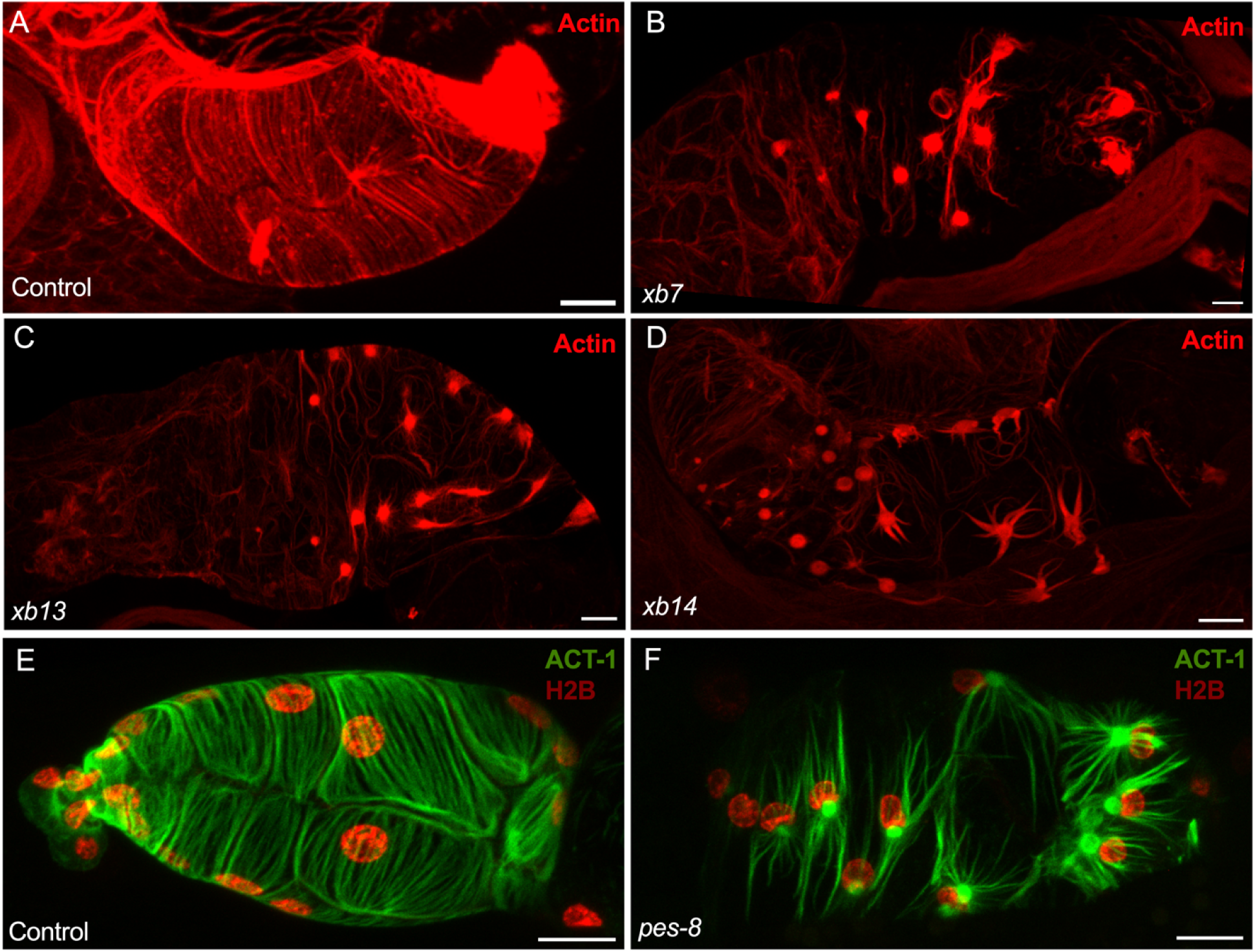
PES-8 anchors actin. A-D) Maximum intensity projections of phalloidin staining shows the occupied spermathecal actin network. A) The actin is organized into long bundles in WT. Phalloidin staining revealed disrupted actin localization in mutants *xb7* (B), *xb13* (C), and *xb14* (D). E and F) Histone/H2b::mCherry and actin/ACT-1::GFP localization in the spermatheca of control (empty vector RNAi), and *pes-8* RNAi knock down animals. Scale bar, 20 μm.

To visualize the dynamics of the actin cytoskeleton during ovulation, we used time lapse confocal microscopy. During the first ovulation in wild-type animals, parallel, aligned actin bundles form (Figure 6A-C, Movie S2) (Kelley et al., 2020). In contrast, in *pes-8* RNAi treated animals, actin bundles appear to detach and become clumped above nuclei of the cells (Figure 6A-F). Before the first ovulation, the *pes-8* spermathecal actin appears similar to wild-type. Once the first oocyte begins to enter the spermatheca (at 15 sec), the actin bundles start to organize. However, by 25 sec, the actin network begins to disassemble, and by 30 sec, the bundles are completely disrupted. By 60 sec, actin aggregates have accumulated around the nuclei, and a complete disruption of the actin structure is observed in both the bag and sp-ut valve regions of the spermatheca (Figure 6D-I) and Movie S3). To enter the uterus, the egg must pass through the spermathecal-uterine (sp-ut) valve. The actin in the sp-ut valve is disrupted, and the valve appears to be distended or ruptured, yet the oocyte does not exit to the uterus, presumably because the spermathecal bag is non-contractile (Figure 6J-M) (Movie S2 and S3).

**Figure 6:**
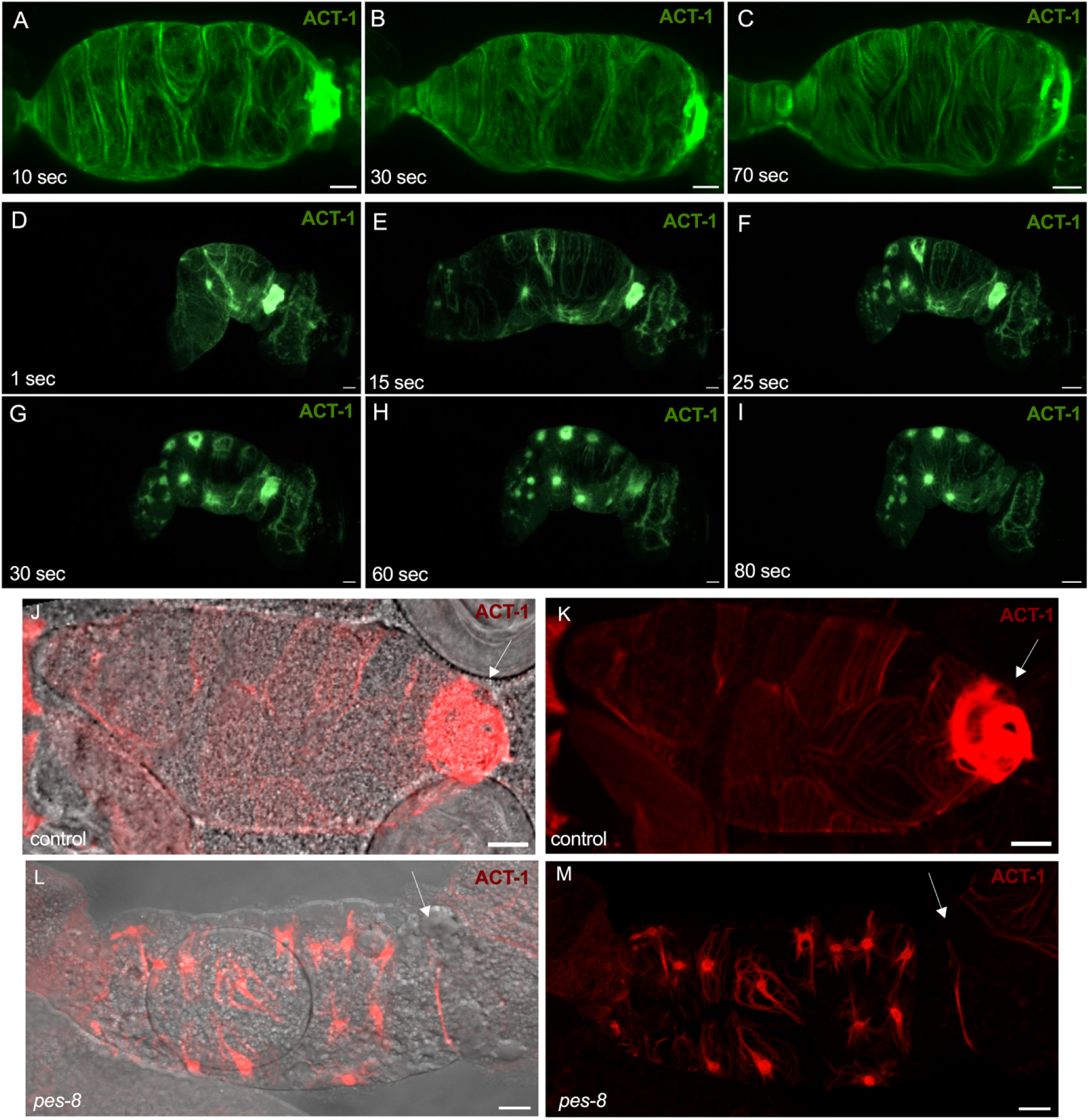
PES-8 anchors actin in the spermathecal bag and valve. A-C) Frames from a confocal time-lapse of actin organization in wild-type from second 10 to second 70 (Movie S2). D-I) Frames from a confocal time-lapse of actin disruption of animals treated with *pes-8* RNAi. Actin bundles in the unoccupied spermatheca before ovulation (D), oocyte entering the spermatheca at second 15 (E), beginning of actin network disruption at second 25 (F) actin disruption at second 30 (G), total aggregation into the nuclei by second 60 (H), complete disruption of actin in the bag and spermathecal-uterine valve (I) (Movie S3). J) Overlay of DIC and phalloidin fluorescent images of actin in the bag and valve in wild-type (control). K) Phalloidin images of actin in the spermathecal bag and valve in control. L) Overlay of DIC and phalloidin fluorescent images of actin with PES-8 depletion. Arrows indicate the valve. M) Phalloidin images of actin in the spermathecal bag and valve with *pes-8* RNAi. Scale bar, 20 μm.

The effect of *pes-8* RNAi on the actin cytoskeleton (Figure 7A and B) reminded us of the phenotype we observed previously while studying the actin crosslinker, FLN-1/filamin (Kelley et al., 2020; Kovacevic & Cram, 2010). Filamin connects transmembrane proteins to the actin network (Calderwood et al., 2001). *fln-1* RNAi produces a similar loss of anchorage of the actin network (Figure 7 B and C). To quantify this effect, we calculated the anisotropy, which gives a measure of alignment of the actin fiber bundles compared to control (Figure 7D). The loss of actin alignment is similar in *pes-8* and *fln-1* RNAi treated animals, similar to the combination of the two RNAis (Figure 7E). We observed similar phenotype treating *pes-8* and *fln-1* mutants with *fln-1* and *pes-8* RNAi, respectively (Figure 7F and G).

**Figure 7:**
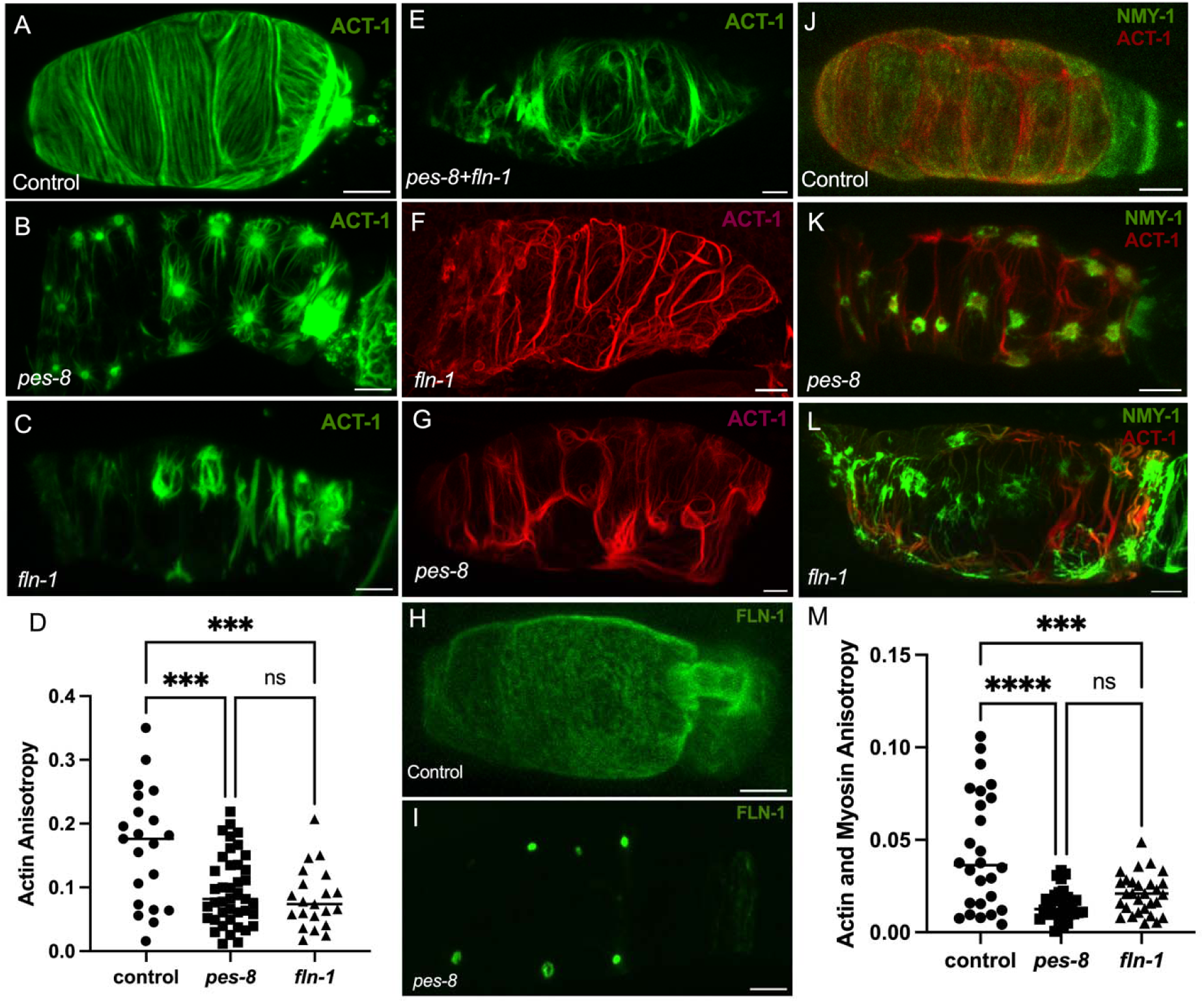
PES-8 is required for anchorage of the actomyosin cytoskeleton and the localization of filamin. A-C) Actin/ACT-1::GFP alignment in the spermatheca of control (empty vector RNAi) (A), *pes-8* RNAi (B), and *fln-1* RNAi (C) knock down animals. D) Actin anisotropy quantification of cells treated with control, *pes-8* and *fln-1* RNAi. E) Actin/ACT-1::GFP alignment in the spermatheca of a 1:1 combination of *pes-8* RNAi and *fln-1* RNAi knock down animals. F) Phalloidin staining of disrupted actin localization in *xb7* mutant with *fln-1* RNAi. G) Actin labeled moeABD::mCherry alignment in the spermatheca of *pes-8* RNAi knock down animals. H and I) Filamin/FLN-1::GFP arrangement in the spermatheca of control (empty vector RNAi) (H), and *pes-8* RNAi (I) knock down animals. J-L) Actin labeled moeABD::mCherry and myosin/NMY-1::GFP bundle alignment in the spermatheca of control (empty vector RNAi) (J), *pes-8* RNAi (K), and *fln-1* RNAi (L) animals. M) Actin and myosin bundle anisotropy of cells treated with control, *pes-8* and *fln-1* RNAis. Scale bar, 20 μm.

Because *pes-8* and *fln-1* RNAi yielded similar actomyosin phenotypes, and filamin anchors actin to transmembrane proteins in other systems (Calderwood et al., 2001), we next asked if PES-8 is required for localization of FLN-1. When PES-8 is depleted in filamin/FLN-1::GFP expressing animals, FLN-1 relocalizes to the region around nuclei rather than decorating actin fiber bundles and expression levels decrease (Figure 7H and I). The mislocalization of FLN-1 may explain the dramatic actin phenotype seen when PES-8 is lost. Conversely, *fln-1* RNAi does not affect the localization of PES-8 (Figure S4B). This suggests PES-8 may work through FLN-1 to anchor or stabilize the actomyosin network on the basal surface of the spermatheca.

Next, we imaged animals expressing NMY-1::GFP and moeABD::mCherry to label non-muscle myosin and filamentous actin, respectively. In wild-type animals, parallel actomyosin bundles are formed on the basal surface of the spermathecal cells (Figure 7J). Depletion of PES-8 results in actomyosin bundle dissociation and aggregation of myosin close to the cell (Figure 7K), similar to the phenotype seen in *fln-1* RNAi (Figure 7L). Actin and myosin anisotropy quantification indicates significant misalignment of the fibers (Figure 7M). Conversely, *nmy-1* RNAi does not affect the localization of PES-8 (Figure S4C).

### PES-8 is required for anchorage of apical junctions

To study whether PES-8 also regulates apical structures, we next examined the localization of the DAC complex by visualizing AJM-1::tagRFP and DLG-1::GFP. Depletion of PES-8 by RNAi results in mislocalization of AJM-1 and DLG-1, indicating PES-8 is required for localization and/or anchorage of these proteins in the cell membrane (Figure 8A-D). Time lapse imaging suggested structures containing AJM-1 and DLG-1 rip apart upon oocyte entry during the first ovulation (Movie S4). Initially, AJM-1 labeled junctions expand upon oocyte entry (Figure 8E-G), but shortly after entry a serial rupture of the junctions occurs (Figure 8H-J). Surprisingly, the oocyte appears to push the AJM-1 labeled junctions to one side of the spermathecal tube (Figure 8K-L). Disruption of actin by *act-1* RNAi or *fln-1* RNAi does not result in the mislocalization of AJM-1 and DLG-1 (Figure S5A, B), suggesting the loss of basal actin does not lead to the mislocalization of DLG-1 and AJM-1. Depletion of DLG-1 and AJM-1 does not disrupt actin bundle formation or anchorage (Figure S6), suggesting the disrupted localization of DLG-1 and AJM-1 is not causing the basal actin phenotype. Additionally, *ajm-1* knockdown does not affect the localization of PES- 8 (Figure S4D). These findings suggest that loss of PES-8 gives rise to disruption of apical junction proteins, potentially independently of its effect on the actin cytoskeleton.

**Figure 8:**
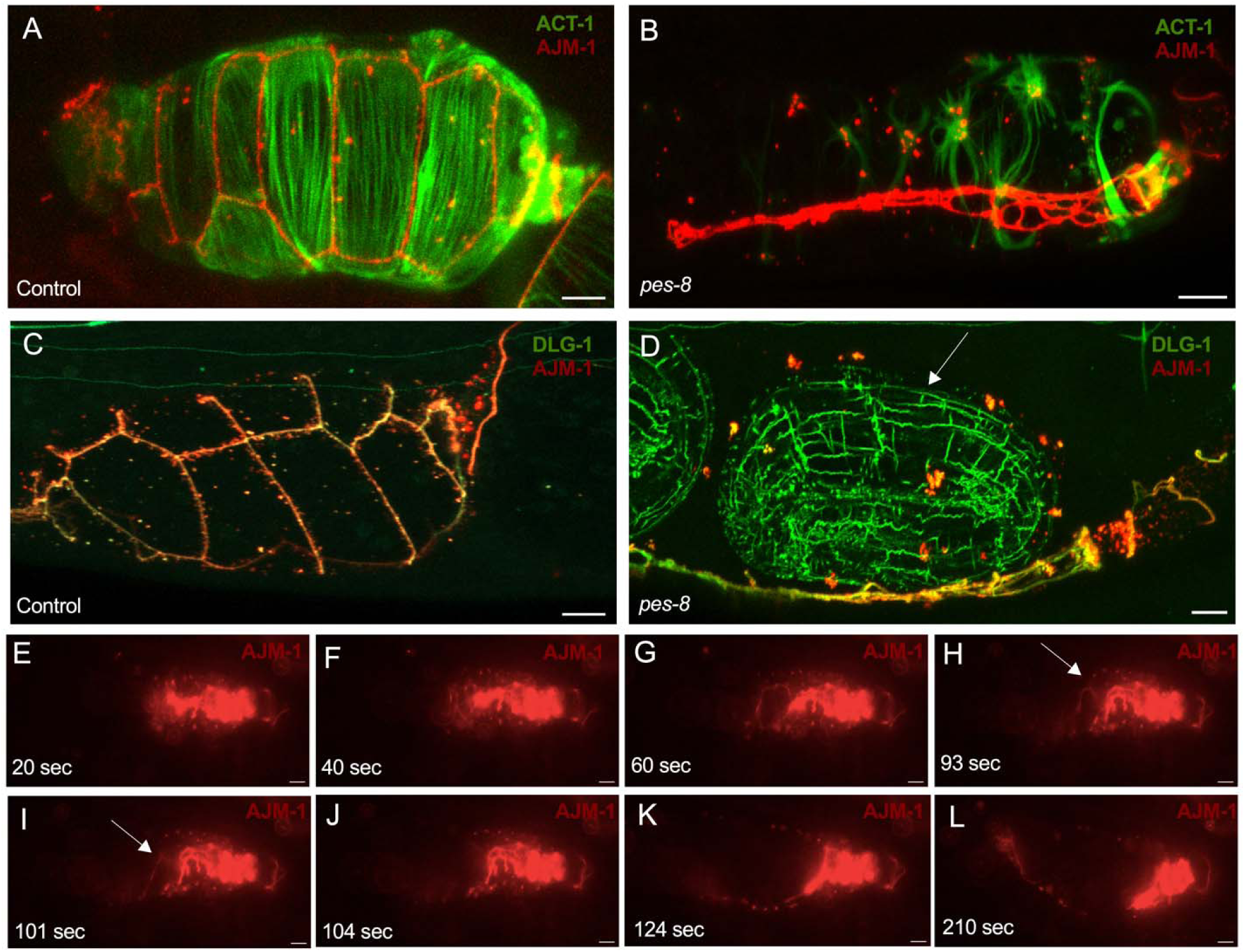
PES-8 is required for the localization of the apical proteins AJM-1 and DLG-1. A and B) ACT-1::GFP and AJM-1::tagRFP expression in the spermatheca of control (empty vector RNAi) (A), *pes-8* RNAi (B) animals. C and D) Discs large/DLG-1::GFP and Apical junction molecule/AJM-1::tagRFP expression in the spermatheca of control (empty vector RNAi) (C), *pes-8* RNAi (D) animals. The arrow indicates DLG-1::GFP expressed in a developing embryo trapped in the spermatheca. Red punctae of AJM-1 around the embryo indicate the embryo remains enclosed in the spermatheca. E-L) Photos of confocal time-lapse of AJM-1 disruption (Movie S4), before oocyte entry (E), upon oocyte entry (F), beginning of expanding the fibers (G), rupture of AJM-1 fiber after oocyte entry (H), rupture of another AJM-1 fiber after oocyte entry (I), J-L) push back of fibers to one side after full oocyte entry. Scale bar, 20 μm.

We next evaluated the localization of HMR-1/E-cadherin, which localizes to the CeAJ (Simske et al., 2003). With *pes-8* RNAi, the localization of HMR-1 is similar to wild-type (Figure 9A-B), and loss of *hmr-1* does not affect the localization of PES-8 (Figure S4E). The cadherin-catenin complex (CCC) is localized apically to the DLG-1/AJM-1 complex (DAC). Therefore, it was surprising that AJM-1 and DLG-1 were disrupted, but HMR-1 was not. A possible explanation is that during the first ovulation (48 hours) when the AJM-1 and DLG-1 are disrupted, HMR-1 is not strongly localized to spermathecal cell junctions. However, it becomes enriched later (56 hrs) in a band at the apical membrane even in *pes-8* depleted animals, suggesting that the cells retain connections with one another and that PES-8 is not required to anchor the cadherin complex (Figure S7).

**Figure 9:**
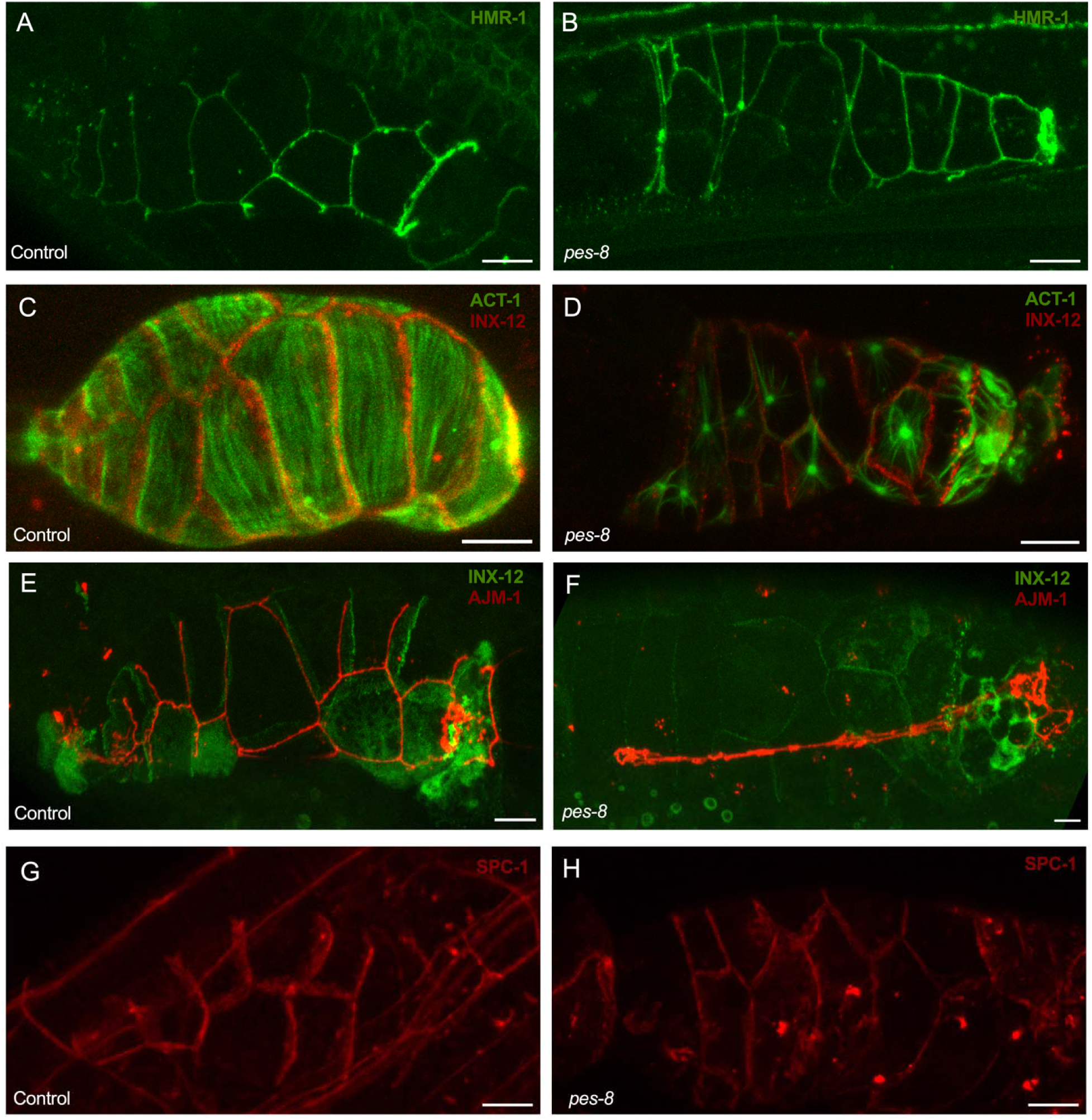
PES-8 is not required for the localization of cadherin, innexin, or spectrin. A and B) Localization of E-Cadherin/HMR-1::GFP in spermatheca of animals treated with control (empty vector RNAi) (A) and *pes-8* RNAi (B). C and D) Innexin /INX-12::mApple and Actin/ACT-1::GFP localization in the spermatheca of control (empty vector RNAi) (C) and *pes-8* RNAi (D) knock down animals. E and F) Innexin/INX-12::GFP and AJM-1::tagRFP expression in the spermatheca of control (empty vector RNAi) (E), *pes-8* RNAi (F) knock down animals. G and H) Localization of Spectrin/SPC-1::mKate2 in the spermatheca of control (empty vector RNAi) (G) and *pes-8* RNAi (H) knock down. Scale bar, 20 μm.

PES-8 is strongly localized to lateral junctions (See Figure 3), therefore, we next tested whether PES-8 is required for the localization of proteins to lateral cell contacts. The localization of the gap junction protein Innexin/INX-12::mApple is not in disrupted *pes-8* RNAi animals, even though the actin cytoskeleton (Figure 9C-D) and AJM-1::tagRFP are mislocalized as expected (Figure 9E-F). We also did not observe any effect on spectrin localization with *pes-8* RNAi treatment (Figure 9G-H). *spc-1* RNAi does not affect the localization of PES-8 (Figure S4 F). These results suggest the integrity of lateral sides of the cells is maintained when PES-8 is disrupted.

### PES-8 partially colocalized with its interactors

To determine whether PES-8 colocalizes with FLN-1, AJM-1 or DLG-1, we made strains expressing green PES-8::GFP coexpressed with FLN-1::RFP, a red actin binding protein (moeABD::mCherry), or AJM-1::tagRFP. We used DLG-1::GFP and AJM-1::tagRFP co-expression as a control for colocalization in the spermathecal bag (Figure 10A). Visual inspection of 3D confocal images and Pearson R correlation analysis indicate only a partial colocalization of PES-8 with FLN-1 and ACT-1 on the basal side (Figure 10B and C), and AJM-1 on the apical side (Figure 10D). This suggests other, unknown, proteins may be mediating the interaction between PES-8, FLN-1 and the apical junction proteins.

**Figure 10:**
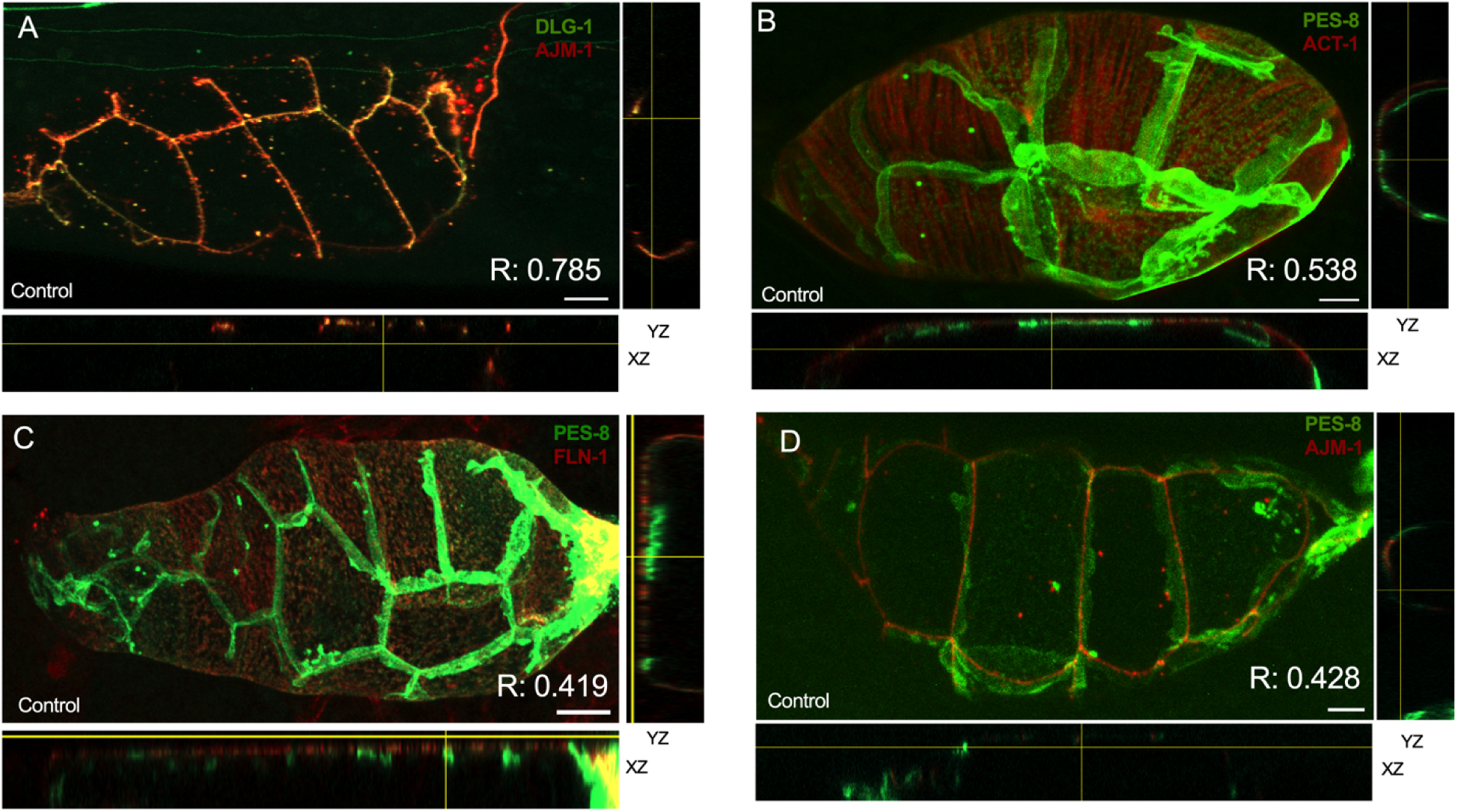
PES-8 partially colocalizes with FLN-1 and AJM-1. Colocalization of PES-8 and actin, filamin and AJM-1. A) DLG-1::GFP and AJM-1::tagRFP expression as control shows high colocalization, Pearson’s R value=0.785. B) PES-8::GFP and moeABD::mCherry expression in the spermatheca of control, *pes-8* RNAi knock down animals, Pearson’s R value=0.538. C) PES-8::GFP and FLN-1::RFP expression in the spermatheca of control, *pes-8* RNAi knock down animals, Pearson’s R value=0.419. D) PES-8::GFP and AJM-1::tagRFP expression in the spermatheca of control, *pes-8* RNAi knock down animals, Pearson’s R value=0.428. Scale bar, 20 μm.

### PES-8 is required for calcium signaling in the spermatheca

Our previous studies highlighted the importance of the PLC-1/phosphatidylinositol signaling in ovulation and oocyte transit in the *C. elegans* spermatheca. PLC-1 generates IP_3_, which initiates Ca^2+^ release from the endoplasmic reticulum via the IP_3_ receptor (Bui & Sternberg, 2002). We explored the impact of silencing PES-8 on Ca^2+^ signaling, by collecting time-lapse fluorescence images of animals expressing GCaMP3. We plotted the data both as 1D traces of total Ca^2+^ levels, and as 2D kymographs with time on the y-axis and space in the x-axis, which allows visualization of the spatial and temporal aspects of Ca^2+^ signaling. Upon oocyte entry in wild-type animals, the sp-ut valve shows a bright pulse of Ca^2+^, which is followed by a quiet period. Once the oocyte is completely enclosed, Ca^2+^ oscillates across the spermatheca, increasing in intensity until peaking concomitantly with distal spermathecal constriction and embryo exit (Figure 11A). To visualize the spatial distribution of the Ca^2+^ pulses over time, normalized traces and kymograms were generated by averaging over the columns of each movie frame for control, *pes-8* RNAi, and *fln-1(tm545)* (Figure 11A-C) (Movies S7-S9). Depletion of *pes-8* and *fln-1* resulted in repetitive pulses of Ca^2+^ in the spermathecal bag and sp-ut valve (Figure 11A-C). Quantification of Ca^2+^ traces indicate a significantly different number of Ca^2+^ peaks between control, *pes-8* RNAi, and *fln-1(tm545)*, reflecting the earlier appearance and persistence of strong pulses as well as the longer duration of the transits (Figure 11D). To visualize and compare multiple *pes-8* RNAi, *fln-1(tm545)* and control transits, we used heat maps to display the intensity of the Ca^2+^ peaks during the transits (Figure 11E-G). To compare the dynamic range of the Ca^2+^ signal during embryo transit, we quantified the half-max (H) of the Ca^2+^ signal (Figure 11H). The dynamic range of the Ca^2+^ pulses in the *pes-8* RNAi treated animals was significantly higher than control, while loss of *fln-*1 results in a lower dynamic range due to a higher baseline. These findings show *pes-8* is required for proper Ca^2+^ regulation in spermatheca. The effect of *pes-8* depletion is not identical to the loss of *fln-1* suggesting other regulators remain to be identified.

**Figure 11:**
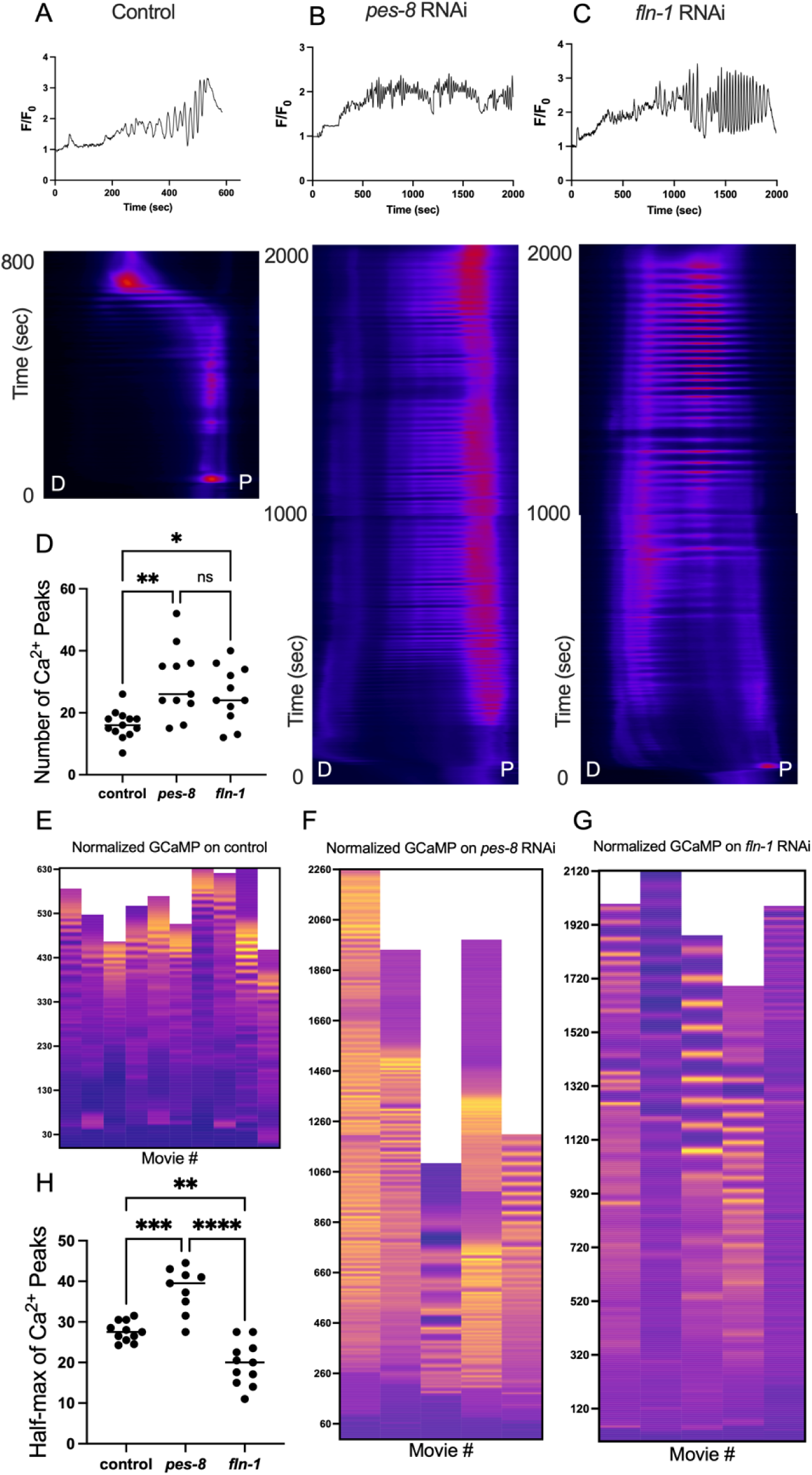
PES-8 regulates Ca^2+^ signaling in the spermatheca. A-C) Normalized Ca^2+^ traces and Ca^2+^ kymographs of control (empty vector RNAi) (A), *pes-8* RNAi (B) and *fln-1(tm545)* (C) in GCaMP expressing animals. D= Distal and P= Proximal spermatheca. D) Ca^2+^ pulse quantification reveals higher number of peaks for *pes-8* and *fln-1(tm545)* compared to control, reflecting the long duration of the embryo transits and the continual Ca^2+^ pulses. E-G) Heat map of the normalized Ca^2+^ traces of control (empty vector RNAi) (n=13) (E), *pes-8* RNAi (n=5) (F) and *fln-1(tm545)* (n=5) (G) Warmer colors indicate stronger Ca^2+^ signal. Traces in A-C correspond to the first column of each heat maps. H) Half-max of the Ca^2+^ signal peaks of control (empty vector RNAi), *pes-8* RNAi and *fln-1(tm545)*.

## Discussion

Here we describe the characterization of PES-8, a novel regulator of contractility. PES-8 does not have significant sequence homology outside of nematodes, however the extracellular domain is predicted to fold into a structure similar to a ZP domain. We show PES-8 is required to anchor the actin cytoskeleton, filamin, and apical junctions against the mechanical strain of oocyte entry. Disruption of these cellular architectures is accompanied by dysregulated calcium signaling, failure of ovulation, and low fertility. Given the putative anchoring function and the predicted ability to form multimers, PES-8 may be involved in both extracellular protein interactions and formation of cytoplasmic signaling complexes.

ZP domains are involved in extracellular protein-protein interactions in sperm-oocyte interactions (Monné & Jovine, 2011) and other contexts (Jovine et al., 2002), and form helical bundles in the apical ECM of many tissues (Drees et al., 2023). Defects in ZP-domain proteins can contribute to infertility, deafness, vascular disorders, inflammatory bowel disease, kidney disease, and cancer (Ghosh & Treisman, 2024). The ZP domain is a protein interaction domain composed of two immunoglobulin-related repeats (Ghosh & Treisman, 2024). Although the overall structure is similar, the extracellular domain of PES-8 is predicted to fold into only one immunoglobulin-like domain. ZP domains are stabilized by disulfide bonds between 5 predicted paired cysteines (Cheng et al., 2005; Cohen et al., 2020), similar to PES-8 which has 15 predicted cysteine bonds and 6 predicted disulfide bonds (Cheng et al., 2005).

Some ZP-domain proteins are anchored to the plasma membrane via a transmembrane domain or GPI linkage. Most are proteolytically cleaved and secreted into the extracellular space (Ghosh & Treisman, 2024). Because PES-8 lacks the predicted furin cleavage site, it is predicted to remain membrane bound. This prediction, in combination with our results that show a requirement for PES-8 in the anchoring and localization of cytoplasmic and membrane proteins, suggests that PES-8 multimers may help organize and stabilize the cells, thereby maintaining integrity and proper cytoskeletal organization during mechanical stress or morphogenesis. This function is similar to other ZP-domain proteins that form extracellular scaffolds to reinforce cell-cell or cell-matrix attachments (Drees et al., 2023), suggesting that PES-8 may play a comparable structural role in the spermatheca. For example, loss of PES-8 results in sp-ut valve rupture, which suggests PES-8 is an important contractility and stability regulator in the sp-ut valve. Future studies will be needed to clarify the role of PES-8 in the sp-ut valve, and to determine whether PES-8 forms a structural lattice or interacts directly with extracellular matrix components to mediate this stabilization.

Our results suggest PES-8 anchors cytoskeletal and membrane proteins, including FLN-1/filamin. On the basal side, disruption of PES-8 leads to mislocalization of FLN-1 and loss of the regular array of actomyosin bundles. Without PES-8, FLN-1, myosin and actin relocalize to the nucleus, likely to the nuclear periphery. This is consistent with our previous results, showing that loss of *fln-1* results in the localization of myosin to the nuclei by the LINC complex components ANC-1 and UNC-84 (Kelley et al., 2020). However, PES-8 and FLN-1 show a low degree of co-localization, therefore, it seems likely another, not yet identified, protein or proteins mediates between PES-8 and FLN-1. PES-8 is also required for the localization and robustness of AJM-1 and DLG-1-containing apical junctions. Without PES-8, these junctions are pushed to the side of the spermatheca. This apical junction phenotype does not seem to be the result of the disorganization of the cytoskeleton at the basal side of the spermatheca, because depletion of neither *act-1* nor *fln-1* results in mislocalization of AJM-1 or DLG-1. These results suggest PES-8 works through different binding partners on the apical and basal sides of the spermathecal tube.

Similar to *C. elegans* embryonic epithelial cells, the spermatheca expresses both the DLG-1/AJM-1 (DAC) and the cadherin-catenin (CCC) complexes, including HMR-1/E-cadherin (Pásti & Labouesse, 2018). These are adhesion complexes found on the apical side of cells that play important roles in cell integrity, cell shape changes, and holding cells together in tissues. The spermatheca lacks tight junctions (Armenti & Nance, 2012) so apical junction complexes are important for tissue integrity. DLG-1 and LET-413/scribble are required for the localization of AJM-1 and the formation of the DAC (Köppen et al., 2001). When PES-8 is lost, AJM-1 and DLG-1 junctions rupture and collapse, suggesting PES-8 may be required not for formation of the junction but to withstand the stress of oocyte entry.

Upon oocyte entry, Ca^2+^ transients spread across the spermatheca and trigger spermathecal contractility. Loss of *pes-8* leads to constant pulses of Ca^2+^ that propagate across the spermathecal bag. Similar abnormal Ca^2+^ signaling is observed when *fln-1* is disrupted (Kovacevic & Cram, 2013). Since loss of *fln-1* disrupts the mitochondrial network and the endoplasmic reticulum as well as the actin cytoskeleton (Kelley et al., 2020), the effect on Ca^2+^ activity is likely to be an indirect result of the loss of cytoplasmic organization and the buffering and storage of Ca^2+^ by these organelles.

PES-8 is expressed not only in the spermatheca but also in other parts of the *C. elegans* reproductive system, including the uterus and vulva, suggesting that this protein may contribute to cytoskeletal organization and Ca^2+^ signaling within these tissues. Such localization suggests a broader functional role for PES-8 beyond ovulation, possibly in coordinating the muscular and signaling activities required for proper egg-laying. Further experiments are needed to determine whether the egg-laying defects observed in *pes-8* mutants arise from dysregulation of uterine or vulval contractility. Additionally, the presence of unfertilized oocytes in ZP-deleted animals suggests PES-8 may play a role in sperm or oocytes.

Our findings indicate that PES-8 is essential for maintaining the structural and functional integrity of the reproductive system, ensuring successful ovulation, fertilization, and egg-laying in *C. elegans*. PES-8 seems to be required for the robustness of the spermatheca to the stress of oocyte entry, as well as for contractility of the spermathecal cells. Many other nematode species have PES-8 homologs, suggesting PES-8 may represent a family of proteins required for tissue robustness and nematode fertility.

## Methods and materials

### Nematode maintenance and strain generation

All animals were grown at 23°C on Nematode Growth Media (NGM) (0.107CM NaCl, 0.25% wt/vol Peptone, 1.7% wt/vol BD Bacto-Agar, 2.5CmM KPO4, 0.5% Nystatin, 0.1CmM CaCl_2_, 0.1CmM MgSO_4_, 0.5% wt/vol cholesterol) and fed OP50 Escherichia coli (Hope, 1999). For experiments, animals were synchronized using an “egg prep,” where gravid hermaphrodites were lysed in an alkaline hypochlorite solution, and washed several times in M9 buffer solution (22CmM KH_2_PO_4_, 42CmM NaHPO_4_, 86 mM NaCl, and 1CmM MgSO_4_) (Hope, 1999). Embryos were then pipetted onto new plates and allowed to grow at 23°C for 52–72Chr, depending on the strain and experiment. All extrachromosomal arrays were injected as described previously (Mello et al., 1991). See the Table 1 for all strains used in this study.

**Table 1:**
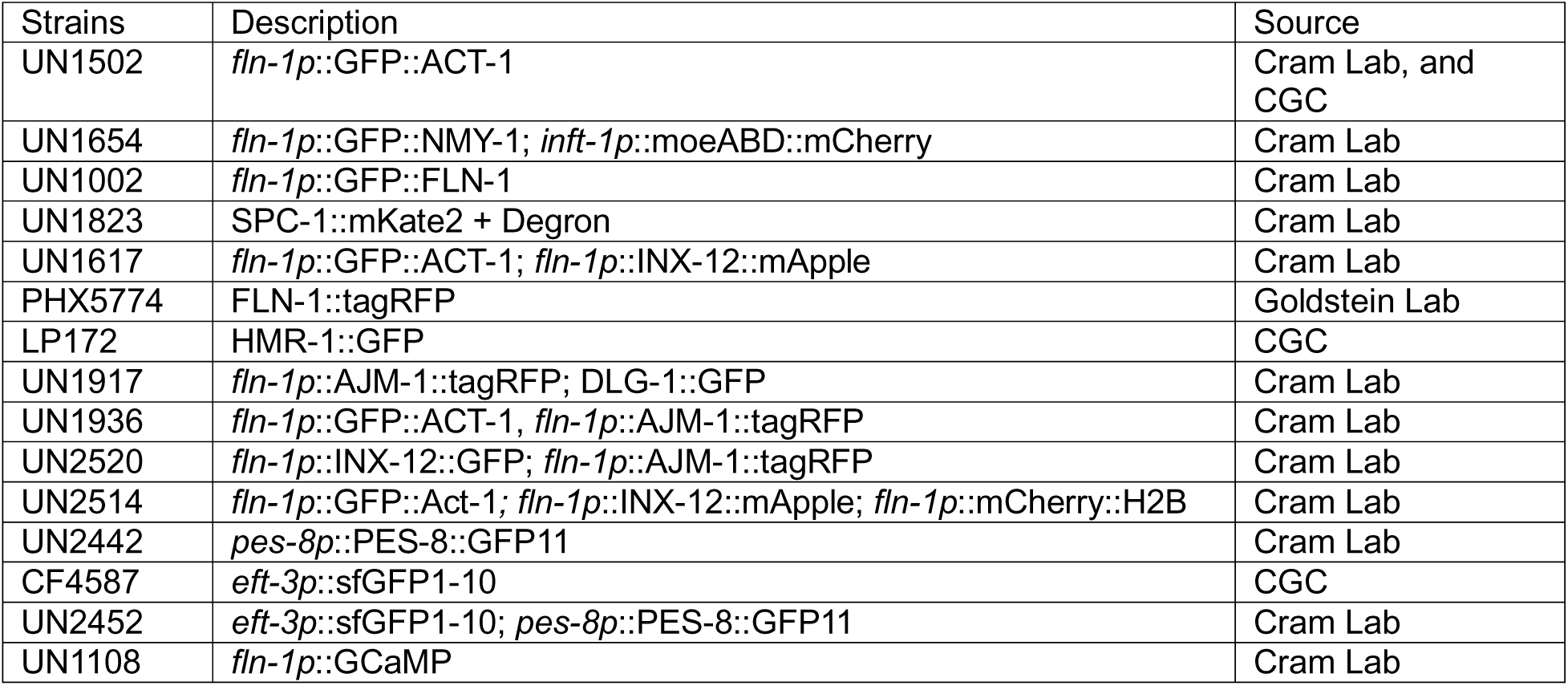
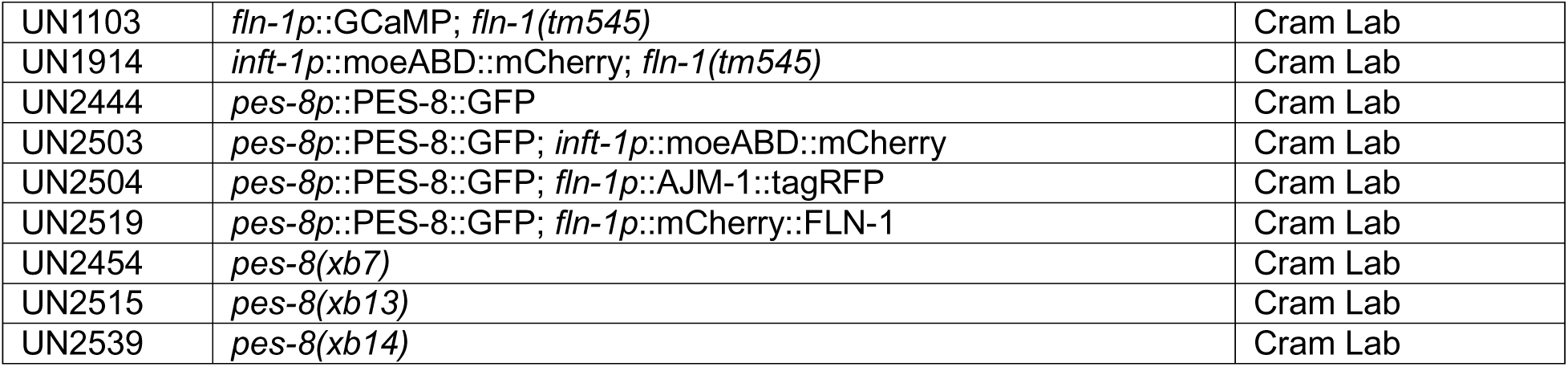
A list of strains.

| Strains | Description | Source |
| --- | --- | --- |
| UN1502 | *fln-1p::GFP::ACT-1* | Cram Lab, and CGC |
| UN1654 | *fln-1p::GFP::NMY-1; inft-1p::moeABD::mCherry* | Cram Lab |
| UN1002 | *fln-1p::GFP::FLN-1* | Cram Lab |
| UN1823 | SPC-1::mKate2 + Degron | Cram Lab |
| UN1617 | *fln-1p::GFP::ACT-1; fln-1p::INX-12::mApple* | Cram Lab |
| PHX5774 | FLN-1::tagRFP | Goldstein Lab |
| LP172 | HMR-1::GFP | CGC |
| UN1917 | *fln-1p::AJM-1::tagRFP; DLG-1::GFP* | Cram Lab |
| UN1936 | *fln-1p::GFP::ACT-1, fln-1p::AJM-1::tagRFP* | Cram Lab |
| UN2520 | *fln-1p::INX-12::GFP; fln-1p::AJM-1::tagRFP* | Cram Lab |
| UN2514 | *fln-1p::GFP::Act-1; fln-1p::INX-12::mApple; fln-1p::mCherry::H2B* | Cram Lab |
| UN2442 | *pes-8p::PES-8::GFP11* | Cram Lab |
| CF4587 | *eft-3p::sfGFP1-10* | CGC |
| UN2452 | *eft-3p::sfGFP1-10; pes-8p::PES-8::GFP11* | Cram Lab |
| UN1108 | *fln-1p::GCaMP* | Cram Lab |
| UN1103 | *fln-1p::GCaMP; fln-1(tm545)* | Cram Lab |
| UN1914 | *inft-1p::moeABD::mCherry; fln-1(tm545)* | Cram Lab |
| UN2444 | *pes-8p::PES-8::GFP* | Cram Lab |
| UN2503 | *pes-8p::PES-8::GFP; inft-1p::moeABD::mCherry* | Cram Lab |
| UN2504 | *pes-8p::PES-8::GFP; fln-1p::AJM-1::tagRFP* | Cram Lab |
| UN2519 | *pes-8p::PES-8::GFP; fln-1p::mCherry::FLN-1* | Cram Lab |
| UN2454 | *pes-8(xb7)* | Cram Lab |
| UN2515 | *pes-8(xb13)* | Cram Lab |
| UN2539 | *pes-8(xb14)* | Cram Lab |

### CRISPR deletion procedure

CRISPR deletion was performed as described (Ghanta et al., 2021). Briefly, synchronized healthy, fed-well, and contamination free N2 worms were obtained by bleaching gravid adults. Adult N2 were injected with 5 μg of Cas9 protein, 0.4 μg/μL of tracrRNA in IDT nuclease free duplex buffer, 0.4 μg/μL crRNA in TE pH 7.5, and 2 ng of a myo-2::mCherry injection marker. Worms were CRISPRed to make *pes-8* alleles of UN2454 (*xb7*), UN2515 (*xb13*), UN2539 (*xb14*). See Table 2 for the crRNA guides used for CRISPR deletion.

**Table 2:**
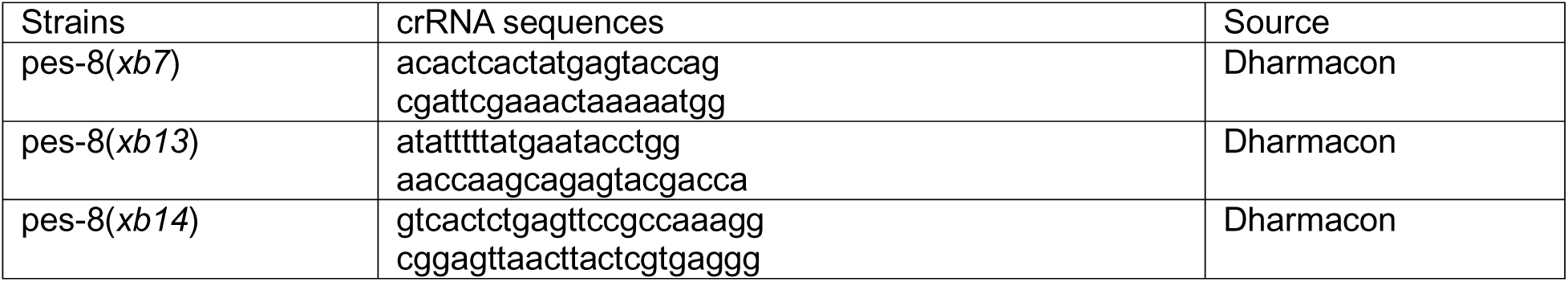
A list of crRNAs.

Primers used for PES-8 cloning and to distinguish *pes-8a* vs *pes-8b* are listed in Table 3.

**Table 3:**
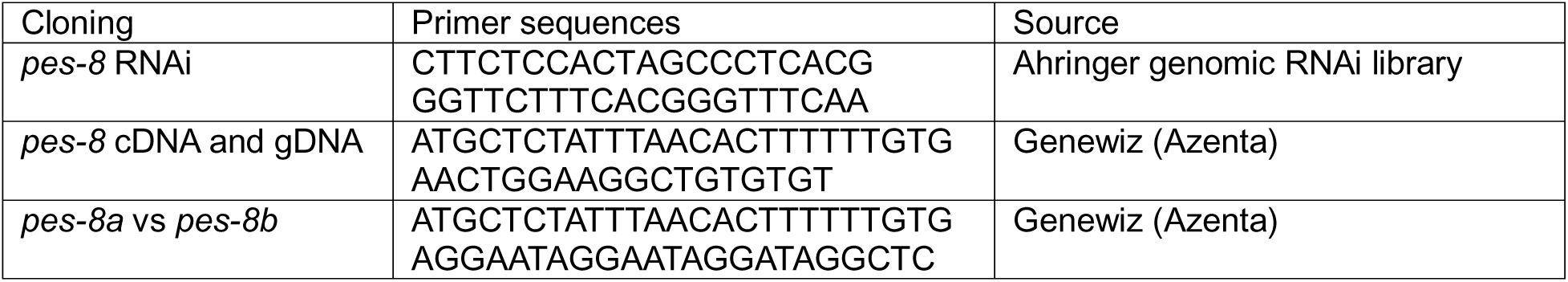
A list of primers.

### RNAi treatment

The RNAi feeding method was used as described previously (Hope, 1999). HT115(DE3) bacteria expressing RNAi constructs in the L4440 backbone were grown shaking overnight in Luria broth at 37°C, then the bacteria were seeded onto NGM plates with 25 μg/ml carbenicillin and 1 mM isopropylthio-β-galactoside (IPTG). Animals were scored for phenotypes as described after propagation at 23°C for 50–56 hours.

### Wide-field Fluorescence and Confocal microscopy image acquisition and processing

Differential Interference Contrast (DIC) and fluorescent timelapse imaging of AJM-1::tagRFP were taken using a 60x oil-immersion objective with Nikon Eclipse fluorescence microscope equipped with a Spot RT3 CCD camera (Diagnostic instruments; Sterling Heights, MI, USA) or Spot RT39M5 with a 0.55x adapter unless otherwise stated. Fluorescence excitation was provided by a Nikon Intensilight C-HGFI 130W mercury lamp and shuttered with a SmartShutter (Sutter Instruments, Novato CA, USA). For acquisition of time-lapse images young adult animals were immobilized with 0.05 micron polystyrene polybeads diluted 1:2 in water (Polysciences Inc., Warrington, PA, USA) and mounted on slides with 5% agarose pads on room temperature. Confocal fluorescence imaging was done on a Zeiss LSM 710 and LSM 800 confocal using Zen software and a Plan-Apochromat 63x/1.40 oil objective lens. Green (GFP) and red (mCherry, tagRFP, mApple) fluorophores were excited with 488 and 561 nm lasers, respectively. Still images were captured at 0.32 μm intervals with each slice being averaged two to four times.

Time lapse GCaMP3 imaging was captured at 1 frame per second, with an exposure time of 75 ms and a gain of 8 for movies obtained with the Spot RT3CCD camera, and with an exposure time of 20 ms and a gain of 8 for movies obtained with the Spot RT39M5. Time-lapse images were only taken of the first 3 ovulations, with preference for the 1st ovulation. The same microscopy image capture parameters were maintained for all imaging. All time-lapse GCaMP3 images were acquired as 1600×1200 pixels for the Spot RT3 OCCD camera or 2448×2048 for the RT39M5 camera and saved as 8-bit tagged image and file format (TIFF) files. All image processing was done using a macro on Image J. All time-lapse images were oriented with the sp-ut valve on the right of the frame and registered to minimize any body movement of the paralyzed animal. An 800×400 region of interest for the Spot RT3 OCCD and 942×471 for the RT39M5 camera encompassing the entire spermatheca was utilized to measure the GCaMP3 signal. The average pixel intensity of each frame was calculated using a custom ImageJ macro. Ca^2+^ pixel intensity (F) was normalized to the average pixel intensity of the first 30 frames prior to the start of ovulation (F_0_) and plotted against time. Data analysis and graphing were performed using MATLAB and GraphPad Prism. MATLAB was used to identify peaks in the GCaMP time series. Local maxima in the time series with a minimum prominence of 0.1 units and a minimum width of 5 units were then identified using the MATLAB ‘findpeaks’ command. These standardized settings were used to analyze all Ca^2+^ traces. Microsoft Excel was used to quantify the amount of time after oocyte entry required to reach either the half the maximum or maximum Ca^2+^ signal as identified by ‘findpeaks’. To determine the rate at which the Ca^2+^ signal changed with time, Kymograms were generated using an ImageJ macro that calculated the average pixel intensity of each column of a frame and condensed it down to one line per frame of the time-lapse image. Every frame of the time-lapse image was stacked to visualize Ca^2+^ dynamics of representative ovulations in both space and time. The half-max value represents the signal intensity halfway between baseline and maximum, serving as a consistent reference point for comparing signal amplitudes across samples. The half-max value was generated by calculating half the difference between the max and the baseline, and adding the baseline (Bouffard et al., 2019). The Fire lookup table color scale was applied to the kymograms using Image J. All MATLAB and ImageJ scripts are available upon request.

### Statistics

Either a Fisher’s exact test (two dimensional x2 analysis) or a one-way ANOVA with a multiple comparison Tukey’s test were conducted using GraphPad Prism on total, dwell and exit transit times of ovulations acquired via time-lapse imaging. For population assays, statistics were performed using the total number of unoccupied spermathecae compared with the sum all other phenotypes. N is the total number of spermathecae counted. For transit phenotype analysis, statistics were performed using the total number of oocytes that exited the spermatheca successfully compared to the sum of all other phenotypes. The Fisher’s exact test was used for both population assays and transit phenotype analysis. Asterisks designate statistical significance (**** p<0.0001, *** p<0.005, ** p<0.01, * p<0.05).

## Abbreviations

CCC: Cadherin–Catenin complex
CeAJs: *C. elegans* apical junctions
DAC: DLG-1/Discs large-AJM-1 complex
GCaMP3: GFP-calmodulin (CaM) and M13 Peptide Version 3
Gf: Gain of function
IP_3_: Inositol trisphosphate
Lf: Loss of function
RNAi: RNA interference
SP-UT: Spermathecal-uterine valve

## Acknowledgements

We thank members of the Cram and Apfeld labs for helpful discussions. Some strains were provided by the CGC, which is funded by NIH Office of Research Infrastructure Programs (P40 OD010440). This work was supported by grants from the National Institutes of Health National Institute of General Medical Sciences (GM110268) and the National Science Foundation – Binational Science Foundation (2419801) to E.J.C.

## Supplementary table and figures

**Table S1:**
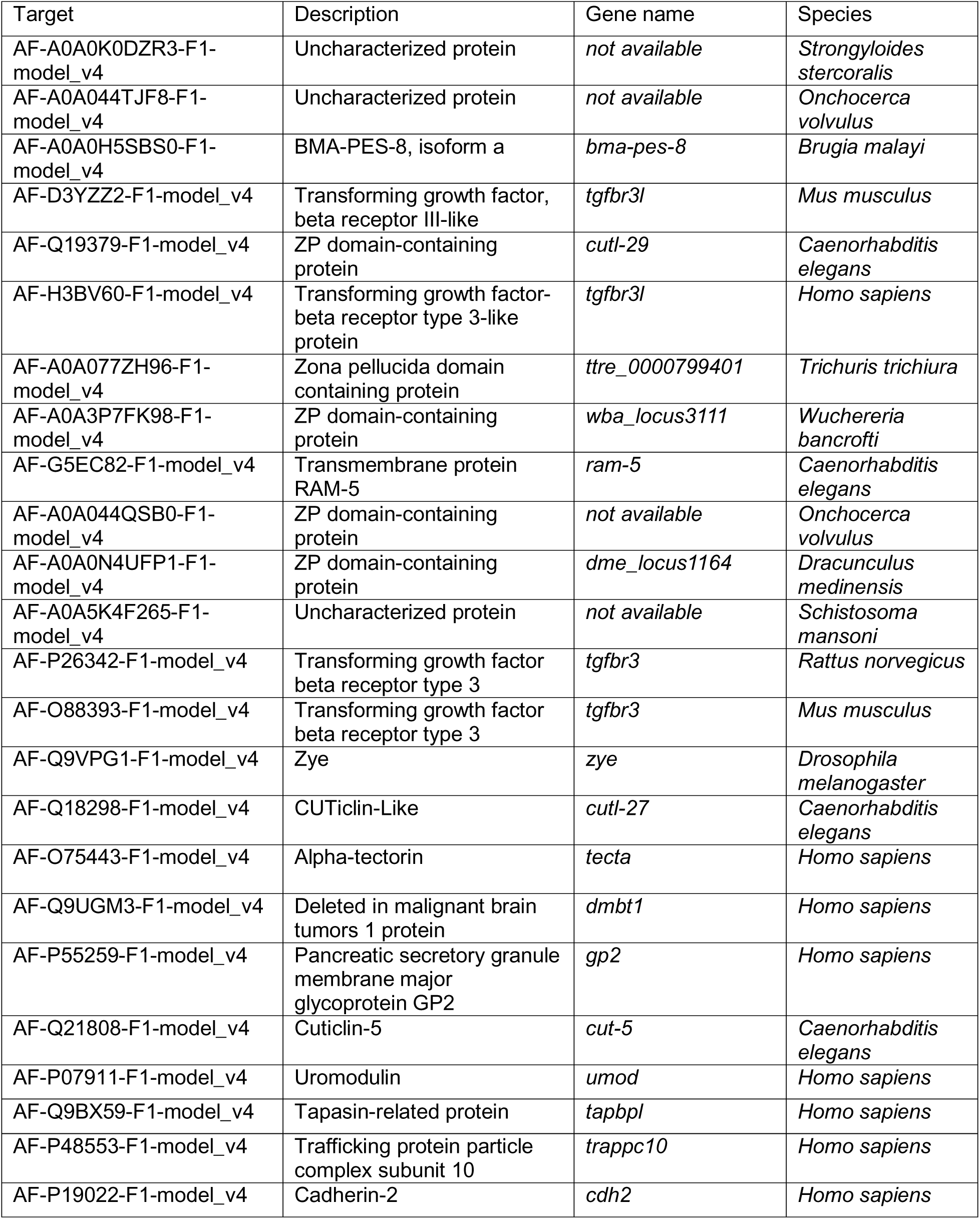
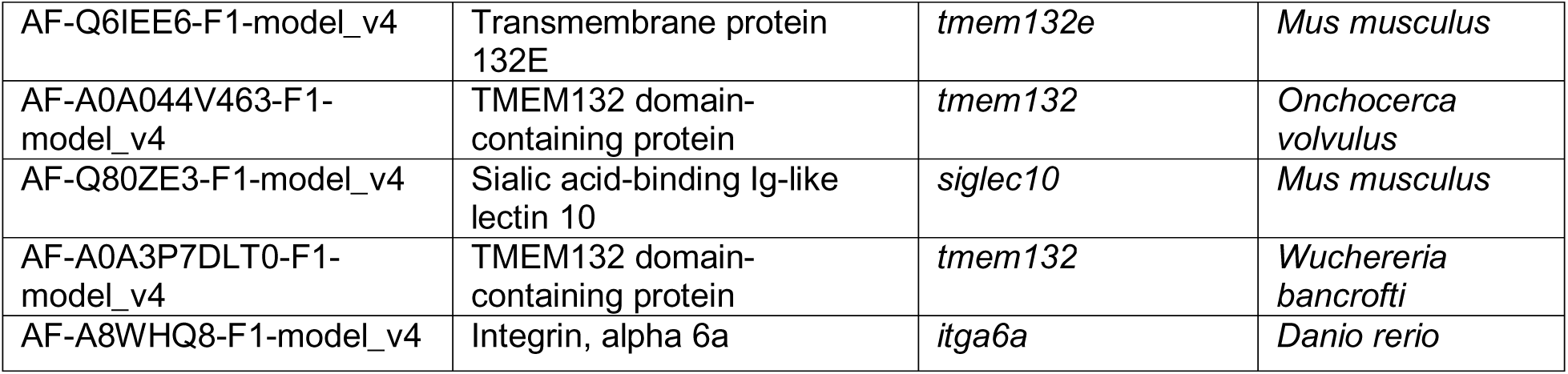
Example proteins with similar folds to PES-8 in other species.

**Figure S1:**
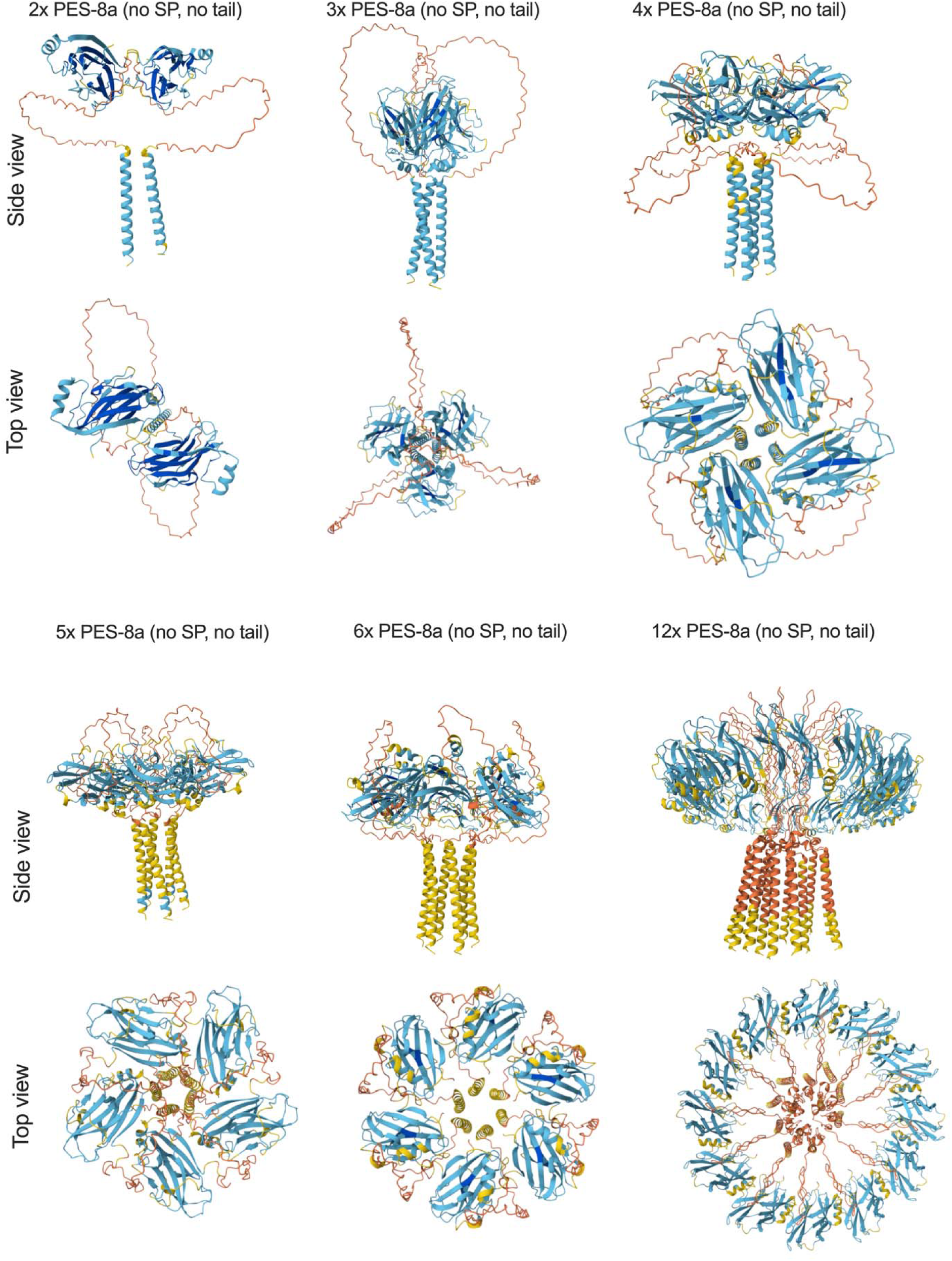
PES-8a AlphaFold predicted multimeric structures.

**Figure S2:**
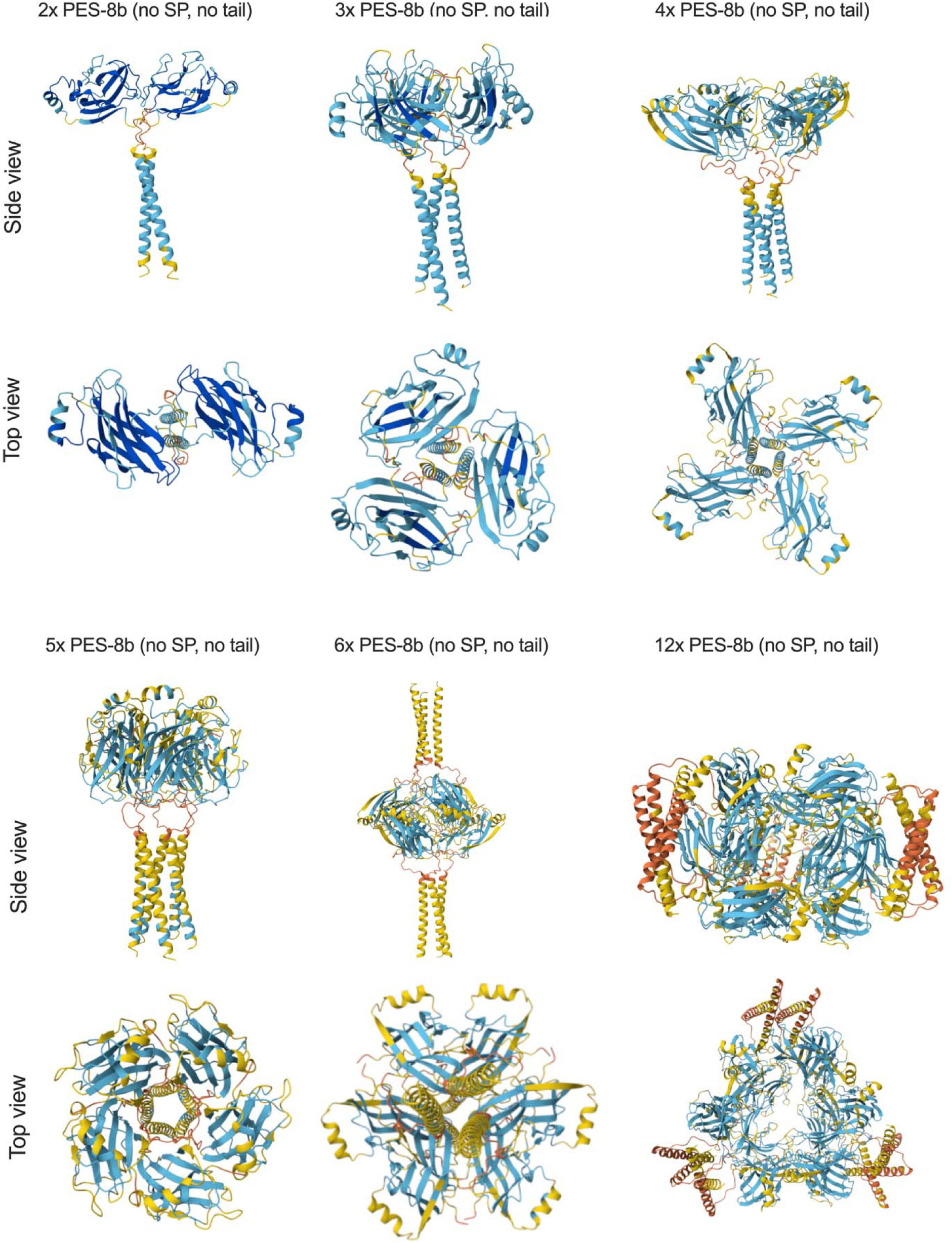
PES-8b AlphaFold predicted multimeric structures.

**Figure S3:**
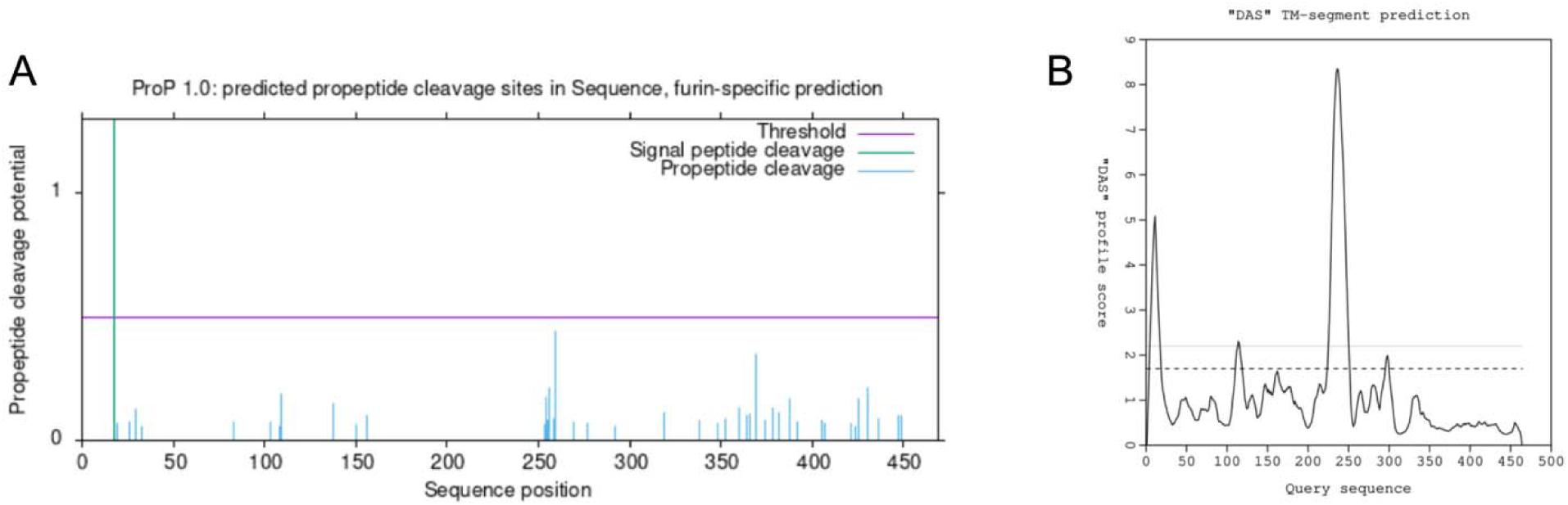
PES-8 membrane sequence prediction. A) ProP 1.0 - DTU Health Tech - Bioinformatic Services (https://services.healthtech.dtu.dk/services/ProP-1.0/) predicts a cleavage site between amino acids 18-19 following the signal sequence and no pro-peptide cleavage site (Duckert et al., 2004). B) DAS - Transmembrane Prediction servers (https://tmdas.bioinfo.se/) predicts two transmembrane domains of PES-8, the C-terminal of which is the putative signal sequence (Cserzö et al., 1997).

**Figure S4:**
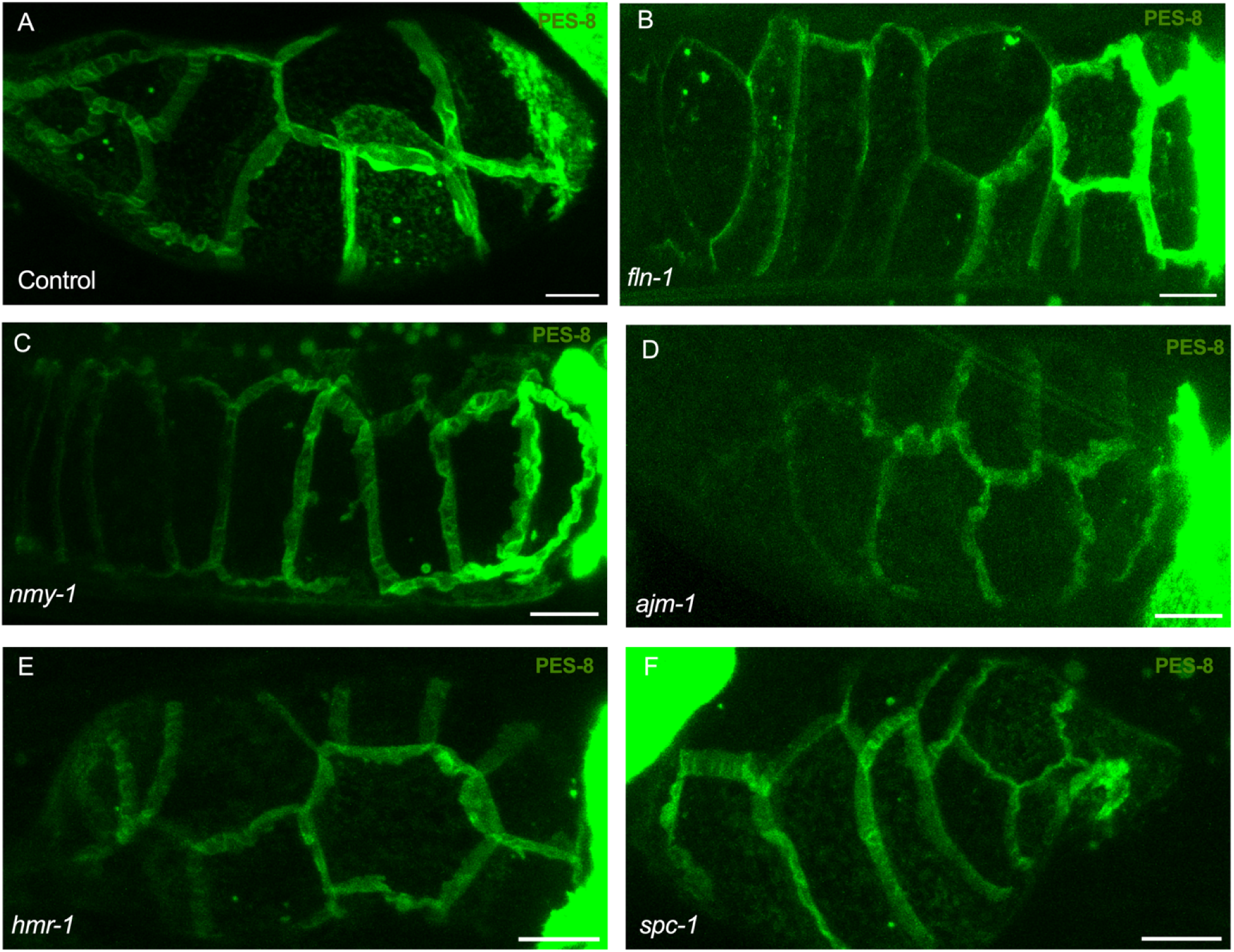
FLN-1, NMY-1, AJM-1, HMR-1 and SPC-1 are not required for the localization of PES-8. Localization of PES-8::GFP in spermatheca with the indicated RNAi treatments does not show significant difference compared to control. Scale bar, 20 μm.

**Figure S5:**
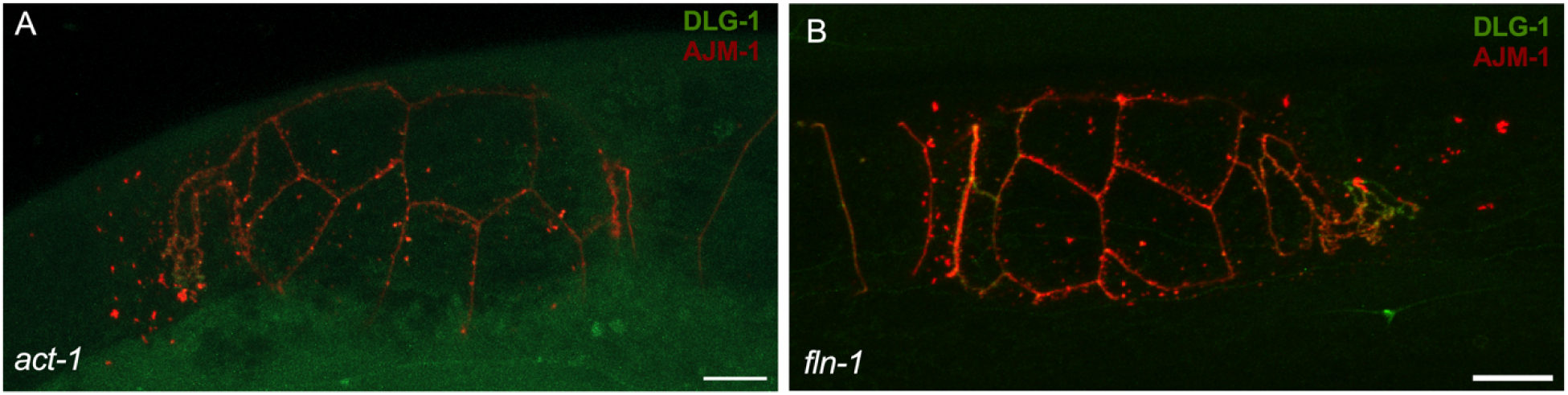
Depletion of ACT-1 and FLN-1 do not affect the localization of DLG-1 and AJM-1. A-B) DLG-1::GFP and AJM-1::tagRFP animals treated with *act-1* RNAi (A) and *fln-1* RNAi (B). Scale bar, 20 μm.

**Figure S6:**
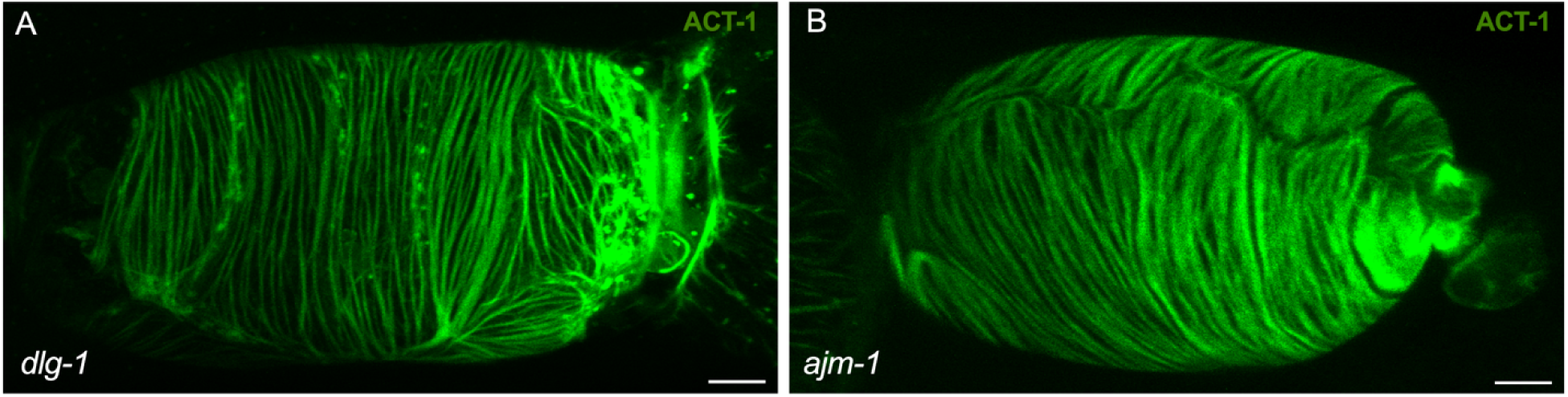
Actin bundles are aligned in DLG-1 and AJM-1 depleted spermathecae: A-B) Alignment of actin bundles in *dlg-1* RNAi (A) and *ajm-1* RNAi (B) knock down conditions do not phenocopy the loss of *pes-8*. Scale bar, 20 μm.

**Figure S7:**
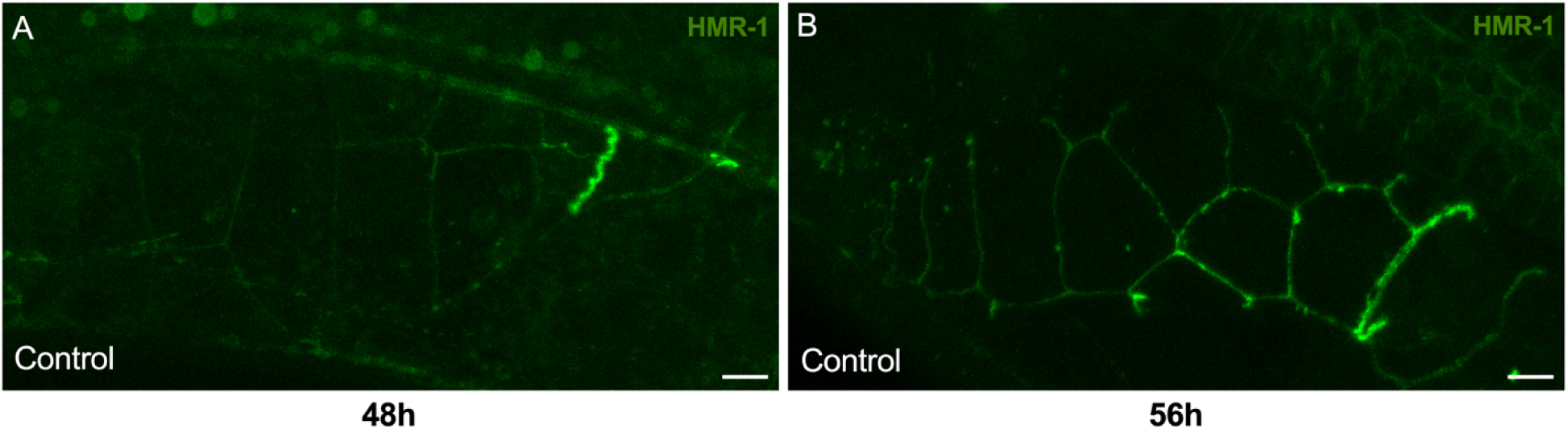
HMR-1 spermathecal expression increases in older animals A-B) Cadherin expression 48h (A) old vs 56h (B) old shows formation of HMR-1 in older animals after a couple of ovulations. Scale bar, 20 μm.

MovieS1: PES-8 expression in spermatheca. Compilation of spermathecal slices from the top (basal) stacks to the cross section of the spermatheca reveals expression of PES-8 on the basal, lateral and apical sides of spermatheca.

MovieS2: Confocal time-lapse of actin organization. Time lapse imaging of ACT-1::GFP expressing animals treated with control RNAi reveals formation of parallel actin bundles in the spermatheca.

MovieS3: Confocal time-lapse of actin disruption. Time lapse imaging of ACT-1::GFP expressing animals treated with *pes-8* RNAi shows actin pulling apart upon oocyte entry and aggregation of the fibers to the nuclei of the cells.

MovieS4: Confocal time-lapse of AJM-1 disruption. Time lapse imaging of AJM-1::tagRFP expressing animals indicates rupture of AJM-1 upon oocyte entry during the first ovulation.

MovieS5: 3D projection of AJM-1 and ACT-1 alignment. AJM-1::tagRFP is localized to apical junctions and and ACT-1::GFP labels parallel bundles in wild-type.

MovieS6: 3D projection of AJM-1 and ACT-1 disruption. AJM-1::tagRFP-containing apical structures are disrupted and pushed to the side and ACT-1::GFP anchorage is lost in *pes-8* RNAi spermathecae.

Movie S7: Ca^2+^ oscillations observed in the spermatheca of animals grown on control RNAi. A wild type ovulation in an animal expressing GCaMP3 shows Ca^2+^ pulses propagating across the spermatheca.

Movie S8: Ca^2+^ oscillations observed in the spermatheca of animals grown on *pes-8* RNAi. The first ovulation of an animal expressing GCaMP3 on *pes-8* RNAi shows constant Ca^2+^ pulses in neck, bag and valve of spermatheca.

Movie S9: Ca^2+^ oscillations observed in the spermatheca of *fln-1(tm545)*. Ovulation of in a *fln-1* null animal expressing GCaMP3 shows constant Ca^2+^ pulses in neck, bag and valve of spermatheca.

