## Supplementary figures and images for "PES-8 is required for Cytoskeletal Organization and Contractility in the *C. elegans* Spermatheca"

### Supplemental Movie 5

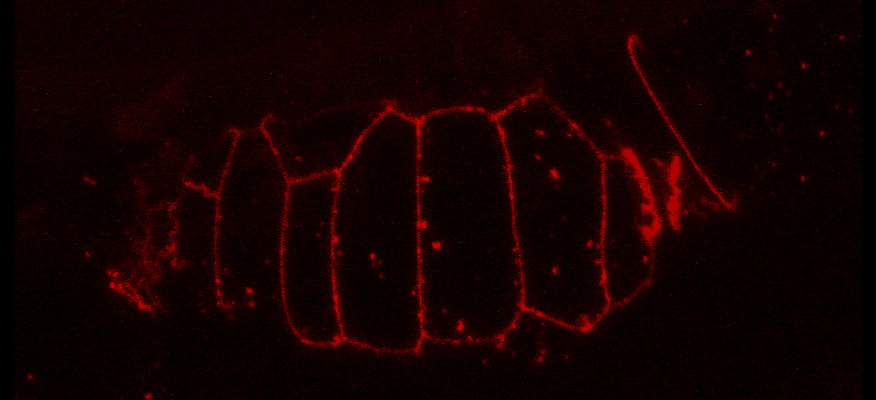
